# Human panepigenome represents epigenomic diversity

**DOI:** 10.64898/2026.08.19.745838

**Authors:** Zheng Dong, Juan F. Macias-Velasco, Xiaoyu Zhuo, Wenjin Zhang, Chad Tomlinson, Juan Jiang, Edward A. Belter, Saifur Rahman Jony, Ronghan Li, Robert S. Fulton, Shihua Dong, Tianjie Liu, Daofeng Li, Human Pangenome Reference Consortium, Ting Wang

**Author notes:** Correspondence to Ting Wang. A complete list of contributing consortium members appears at the end of this paper (Human Pangenome Reference Consortium).

## Abstract

The human pangenome captures genetic diversity beyond a single linear reference, yet conventional DNA methylome maps largely assume that the underlying CpG substrate is fixed. Here we present a first draft of the human panepigenome on DNA methylation, generated from 440 haplotype-resolved long-read methylomes spanning 26 globally distributed populations. This map jointly captures CpG presence, absence and methylation across diverse human haplotypes, revealing 12.8 million CpGs absent from GRCh38 and an average of 1.2 million variant-associated CpGs (var-CpGs) per methylome. Var-CpGs captured population-associated epigenomic variation beyond that represented by shared reference CpGs, and structural variants could create or remove entire CpG islands, frequently through mobile element insertions. Promoter var-CpG methylation was associated with transcript expression, with most associations persisting after adjustment for the underlying CpG-altering variant, whereas a subset showed evidence of mediation or variant-by-methylation interactions. CpG-altering variants were also enriched among molecular QTLs in HPRC2 and across GTEx tissues, and among clinically and pharmacogenomically annotated loci. Together, these results establish a haplotype-resolved framework in which human epigenomic diversity reflects not only variation in methylation level but also genetic variation in presence or absence of CpG substrates, providing a foundation for panepigenomic studies across tissues, populations and disease contexts.

## Main

A pangenome moves beyond a single linear reference by representing genetic diversity across populations, including non-reference and structurally divergent sequences^1–3^. Pangenome resources have improved variant discovery and the representation of genomic regions poorly captured by conventional references, providing a foundation for interpreting genetic and functional variation^4^. However, functional genomic maps remain largely anchored to linear reference coordinates, leaving non-reference and structurally variable regions incompletely characterized. This limitation is especially important for DNA methylation (DNAm), because genetic variation can alter not only methylation levels at existing CpGs but also whether a CpG is present at all. As DNAm contributes to gene regulation, development, aging and disease^5–12^, excluding genetically variable CpGs leaves an important dimension of human regulatory diversity unresolved. Just as reference epigenome projects extended the Human Genome Project by mapping regulatory states across cell types and tissues^13^, a human panepigenome would extend the pangenome concept by cataloging CpG presence, absence and methylation across diverse genomes (Fig. 1a). This map will connect genetic variation encoded in the human pangenome with population-scale epigenetic and regulatory variation.

**Fig. 1:**
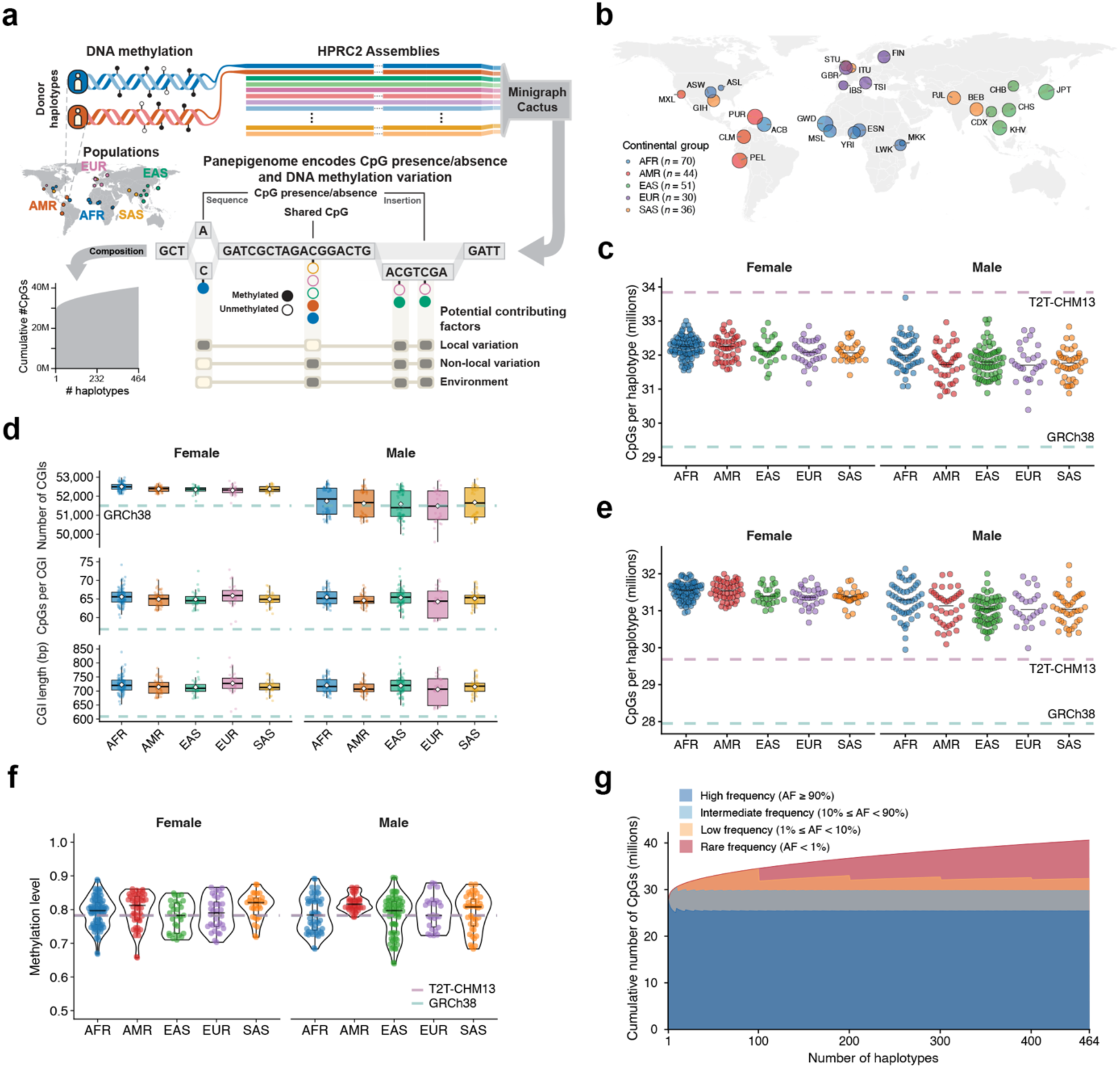
Haplotype-resolved human methylomes and the human panepigenome. **a**, Conceptual overview of the human panepigenome. The human panepigenome integrates haplotype-resolved genome sequences with long-read DNA methylation profiles to capture both CpG presence or absence and methylation variation across genetically diverse haplotypes. Unlike conventional reference-based methylome maps, which measure methylation only at CpGs represented in a single linear reference genome, the panepigenome identifies non-reference CpGs and variant-associated CpGs (var-CpGs) generated or disrupted by single-nucleotide variants (SNVs), indels and SVs, thereby expanding the measurable human epigenomic landscape. Graph-based, single-base-resolution coordinates enable epigenetic variation, including CpG gain or loss and methylation variation, to be linked to local variation (i.e., genetic variants that generate or disrupt CpG sites), nearby genetic variation, and environmental factors, providing a population-scale framework for analyzing epigenomic variation beyond the linear reference genome. Three-letter labels indicate continental groups. AFR, African; AMR, American; EAS, East Asian; EUR, European; SAS, South Asian. **b**, Summary of the HPRC2 sample continental groups and populations on a map of Earth as predominantly defined by the 1000 Genomes Project, colored by continental groups. HG002 and HG005 were excluded. Three-letter labels indicate human populations. ACB, African Caribbean in Barbados; ASL, African Americans living in St. Louis, Missouri; ASW, African Ancestry in Southwest US; BEB, Bengali in Bangladesh; CDX, Chinese Dai in Xishuangbanna; CHB, Han Chinese in Beijing, China; CHS, Han Chinese South; CLM, Colombian in Medellin, Colombia; ESN, Esan in Nigeria; FIN, Finnish in Finland; GBR, British from England and Scotland; GIH, Gujarati Indians in Houston, Texas, USA; GWD, Gambian in Western Division; IBS, Iberian Populations in Spain; ITU, Indian Telugu in the UK; JPT, Japanese in Tokyo, Japan; KHV, Kinh in Ho Chi Minh City, Vietnam; LWK, Luhya in Webuye, Kenya; MKK, Maasai in Kinyawa, Kenya; MSL, Mende in Sierra Leone; MXL, Mexican Ancestry in Los Angeles, California, USA; PEL, Peruvian in Lima, Peru; PJL, Punjabi in Lahore, Pakistan; PUR, Puerto Rican in Puerto Rico; STU, Sri Lankan Tamil in the UK; TSI, Toscani in Italia; YRI, Yoruba in Ibadan, Nigeria. Of note, ASL is not a population defined by the 1000 Genomes Project and is represented by only one sample. **c**, Number of CpGs in 462 haplotype-resolved assemblies, GRCh38 and T2T-CHM13. Each haplotype-resolved assembly is shown as a dot; teal and purple dashed horizontal lines indicate GRCh38 and T2T-CHM13, respectively. Haplotype-resolved assemblies are grouped by sex and shown separately for male and female individuals. **d**, Haplotype-resolved CpG island (CGI) annotations across continental groups, stratified by sex. Box plots show the distributions of the number of CGIs, mean number of CpGs per CGI and mean CGI length across haplotype-resolved assemblies. Points represent individual haplotype-resolved assemblies and are colored by continental group; white diamonds indicate group means. Teal dashed horizontal lines indicate the corresponding GRCh38 reference values. haplotype-resolved assemblies are grouped by sex and shown separately for male and female individuals. **e**, Comparison of the number of CpGs with methylation profiles across 462 haplotype-resolved assemblies, GRCh38 and T2T-CHM13. CpGs were called separately after alignment to each individual haplotype-resolved assembly, GRCh38, or T2T-CHM13. Each haplotype-resolved assembly is shown as a dot; teal and purple dashed horizontal lines indicate GRCh38 and T2T-CHM13, respectively. Haplotype-resolved assemblies are grouped by sex and shown separately for male and female individuals. **f**, Distribution of genome-wide median CpG methylation levels across continental groups, stratified by sex. Points represent individual haplotype-resolved assemblies and are colored by continental group; violin plots show the distribution within each group. Black horizontal marks indicate group medians. Teal and purple dashed horizontal lines indicate the median methylation levels obtained after alignment to T2T-CHM13 (78.3%) and GRCh38 (78.2%), respectively. **g**, Growth curve showing cumulative CpG discovery across haplotype-resolved assemblies, stratified by CpG presence frequency. The panepigenome includes CpGs from 462 haplotype-resolved assemblies, GRCh38 and T2T-CHM13, represented using the HPRC2 Minigraph-Cactus graph with GRCh38 as the backbone. CpGs were stratified by presence frequency into bins of <1%, ≥1% to <10%, ≥10% to <90% and ≥90%.

Population-scale DNAm studies reveal human epigenomic diversity but remain restricted to CpGs represented in a single linear reference genome^14–22^. Conventional reference-based approaches therefore cannot capture non-reference CpGs, including those created or removed by structural variants (SVs), or resolve methylation across alternative haplotypes^23–26^. Pangenome graphs provide a coordinate system for representing genetic diversity at single-nucleotide resolution^27–29^, enabling CpGs and their methylation levels to be analyzed across reference and non-reference genomic contexts.

The Human Pangenome Reference Consortium Release 2 (HPRC2) expands HPRC1, with long-read data from 231 individuals across 26 populations^30^. Long-read sequencing and assembly enable haplotype-resolved genome representation, while sequencing native DNA molecules retains endogenous base-modification signals^31–33^. Together, these features enable joint analysis of genetic variation and DNAm on haplotypes, including CpG gain and loss and methylation variation within structurally complex regions^33–35^, providing a basis for population-scale panepigenomic mapping.

Here, we leverage 440 haplotype-resolved PacBio HiFi methylomes from lymphoblastoid cell lines (LCLs) of 220 individuals, predominantly from the 1000 Genomes Project (1KG) cohort^36^, together with near-telomere-to-telomere assemblies and HPRC2 graph coordinates, to construct a first draft of the human DNAm panepigenome. Focusing on 5-methylcytosine (5mC), we identify 12.8 million CpGs absent from GRCh38 and capture epigenomic variation across three layers: methylation variation at shared CpGs, CpG gain or loss at var-CpGs, and an integrated signal combining CpG presence or absence with methylation on CpG-containing haplotypes. Var-CpGs capture widespread population-associated epigenomic variation, SVs can create or remove entire CpG islands (CGIs), and var-CpG methylation is associated with transcript expression. CpG-altering variants are also enriched among molecular quantitative trait loci (QTLs) and clinically or pharmacogenomically annotated loci. Although demonstrated only in LCLs, the framework is extensible to other tissues and cell types. Together, this work establishes a population-scale, haplotype-resolved framework for understanding how genetic diversity shapes CpG presence, absence and methylation, and how multiple dimensions of epigenomic variation relate to population diversity, genome regulation and clinically annotated loci.

### Haplotype-resolved human panepigenome

A complete representation of human epigenomic diversity requires resolving not only methylation variation, but also variation in the CpG landscape itself across genetically diverse haplotypes.

#### Global long-read haplotype-resolved LCL methylomes

To capture epigenomic variation across human genetic diversity, we analyzed 231 LCL samples from HPRC2, representing 26 populations across five continental groups: Africa (AFR, *n* = 70), the Americas (AMR, *n* = 44), East Asia (EAS, *n* = 51), Europe (EUR, *n* = 30), and South Asia (SAS, *n* = 36)^30^ (Fig. 1b and Supplementary Table 1). CpG landscapes were characterized across 462 near-telomere-to-telomere (near-T2T) haplotype-resolved genome assemblies generated by HPRC2.

Across these assemblies, we identified an average of 32.0 million CpGs per haplotype, exceeding GRCh38 (29.3 million) but remaining below T2T-CHM13 (33.8 million) (Fig. 1c and Supplementary Table 2). After accounting for assembly size, CpG density remained higher than in GRCh38 but slightly lower than in T2T-CHM13, consistent with improved representation of repetitive regions, including pericentromeric satellites and segmental duplications (Extended Data Fig. 1a,b). The expanded CpG repertoire also altered annotation of CpG-rich regulatory elements. Compared with GRCh38, haplotype-resolved assemblies contained an average of 516 additional CGIs per methylome (Fig. 1d).

Long-read sequencing enables DNAm to be resolved at the haplotype level^37^. After DNAm quality control, 220 of 231 individuals were retained for haplotype-resolved methylation profiling, with a mean genomic coverage of 58.4× (Extended Data Fig. 2 and Supplementary Table 3). Based on alignments, each methylome captured an average of 31.3 million CpGs, ∼3.3 million and ∼1.6 million more than GRCh38-and T2T-CHM13-based alignments, respectively, highlighting the value of personalized genome references for epigenomic profiling (Fig. 1e).

Genome-wide, haplotype-resolved methylomes had a median DNAm level of 80.3%, slightly higher than estimates from GRCh38- and T2T-CHM13-based alignments (78.2% and 78.3%, respectively; Fig. 1f and Supplementary Table 4). To facilitate comparison with existing clinical and research resources^38^, GRCh38 was used as the reference for downstream analyses. The higher methylation was largely attributable to non-reference CpGs, which were more highly methylated and less frequently hypomethylated than reference CpGs (Extended Data Fig. 3a,b). Non-reference CpGs were also unexpectedly more highly methylated than reference CpGs within CGIs and promoters, revealing distinct methylation landscapes in CpG-rich regulatory regions in LCLs (Extended Data Fig. 3c).

#### Diversity across haplotype-resolved human methylomes

Haplotype-resolved methylomes enabled population-scale analysis of epigenomic diversity across human haplotypes. Consistent with greater genetic diversity in AFR populations^36,39–44^, AFR assemblies contained the most CpGs in both females and males, as well as the most CGIs (Fig. 1c,d). Non-reference CpG enrichment relative to reference CpGs also varied across continental groups (Extended Data Fig. 1b). Although non-reference CpGs were depleted from gene bodies in all groups, this depletion was weaker in AFR, suggesting population-associated differences in their genomic distribution (Supplementary Table 5).

AMR methylomes showed the greatest between-population variability within continental groups in assembly-driven CpG counts, consistent with the complex demographic history of admixed American populations^45^ (Supplementary Table 6). This variability was particularly pronounced in males and produced the largest sex difference in CpG counts among continental groups (Extended Data Fig. 4a,b). These patterns may reflect sex-associated differences in ancestry composition, as reported for genetic variants^46^, or sampling variation; larger cohorts will be needed to distinguish these possibilities.

Genome-wide DNAm levels also varied across continental groups and sexes, with distinct patterns of between-population variability (Extended Data Fig. 4c and Supplementary Table 6). These patterns differed from variation in CpG counts and previously reported population differences in genetic variant counts^44^, suggesting that genome-wide methylation diversity is not explained by genetic diversity alone and may also reflect environmental or stochastic influences^15–19^.

Non-reference CpG methylation further varied across continental groups both genome-wide and across multiple genomic elements (Extended Data Fig. 3a,c). Together, these findings reveal substantial haplotype-scale methylome diversity and identify non-reference CpGs as an important source of population-associated methylation variation.

#### Draft graph-based human panepigenome

Haplotype-resolved methylomes enabled construction of a draft human panepigenome that captured population-scale variation in CpG presence, absence and methylation at single-nucleotide resolution. We integrated these methylomes into the HPRC2 Minigraph-Cactus graph using GRCh38 as the backbone. The resulting panepigenome comprised 40.6 million CpGs, including 8.2 million rare (<1%), 2.6 million low-frequency (1–10%), 4.3 million intermediate-frequency (10–90%) and 25.4 million high-frequency (≥90%) CpGs, of which 12.8 million were absent from GRCh38 (Fig. 1g).

Notably, 130.7K high-frequency non-reference CpGs were absent from GRCh38 despite being broadly represented across sampled haplotypes. Conversely, 265.1K CpGs present in GRCh38 were rare or low-frequency across HPRC2 haplotypes, indicating that some reference CpGs are absent from most sampled haplotypes. Thus, the graph-based panepigenome captures CpGs missing from GRCh38 while identifying reference CpGs that are not broadly represented across human genomes, helping to address one source of reference bias in population-scale methylome analyses.

### Genetic variation shapes the methylation landscape

Because genetic variation can create or remove CpGs, differences in DNA sequence directly shape the substrate on which methylation occurs.

#### CpG gain and loss shaped by genetic variants

We define CpGs gained or lost through genetic variation as variant-associated CpGs (var-CpGs; Fig. 2a). To characterize var-CpGs, we intersected assembly-derived CpGs with single-nucleotide variants (SNVs), indels and SVs identified relative to GRCh38 using alignment- and assembly-based approaches, leveraging long reads to improve variant detection in complex regions^30,47,48^. Each methylome contained an average of ∼1.2 million var-CpGs, comprising 629.0K gains and 546.8K losses (Fig. 2b and Supplementary Table 7). Across methylomes, the median genome-wide DNAm level at var-CpGs was 81.8%, and their methylation distributions differed from those of CpGs that did not overlap genetic variants (Fig. 2c,d and Supplementary Table 8).

**Fig. 2:**
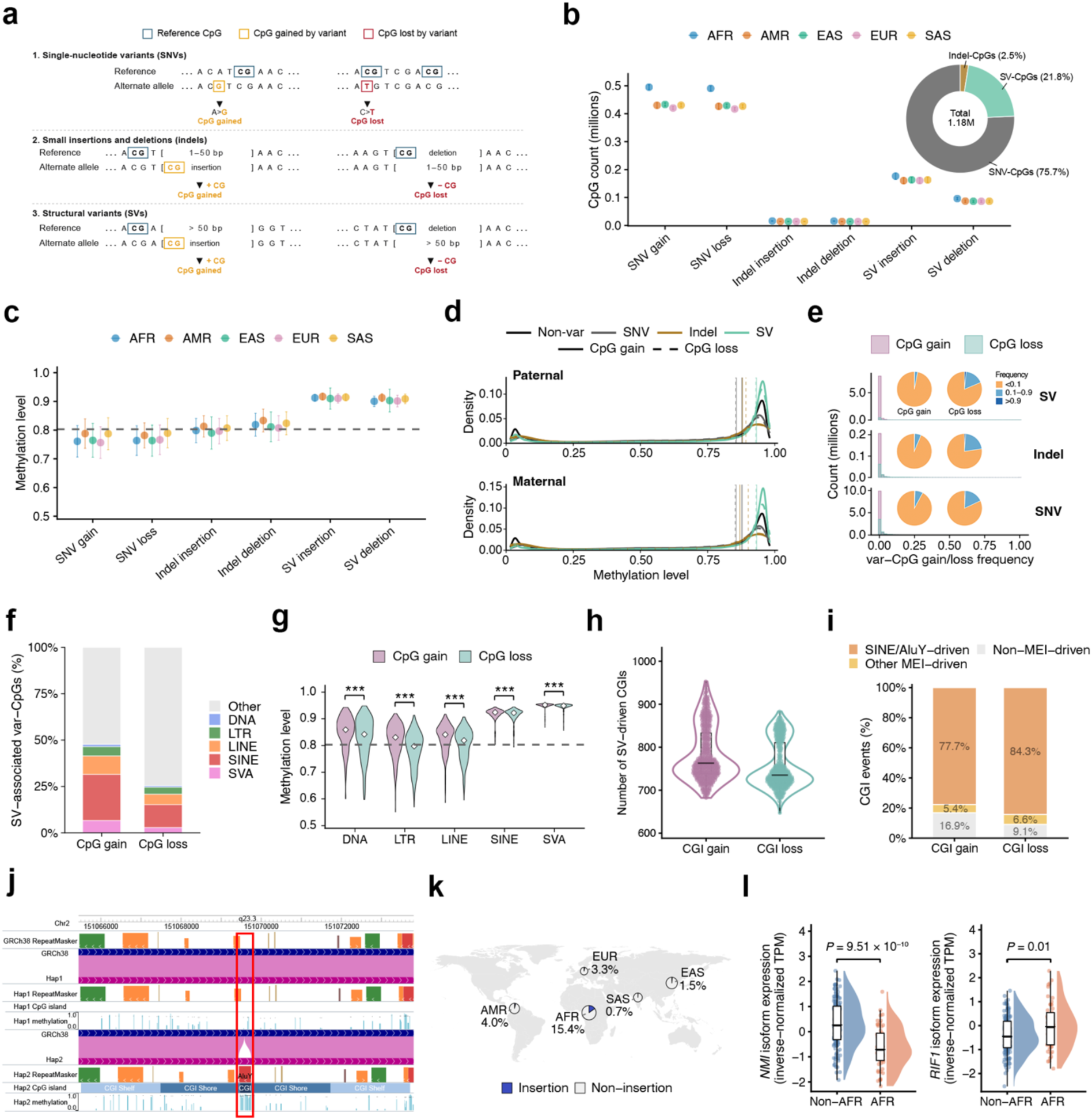
Genetic variation shapes human methylomes. **a**, Schematic illustration of CpG gain and loss caused by genetic variants. CpG dinucleotides are shown on the forward strand for illustration. Variants can create (gain) or abolish (loss) CpG sites. **b**, CpG gain and loss across genetic variant classes and continental groups. Points indicate the median number of CpGs associated with SNV gain, SNV loss, indel insertion, indel deletion, SV insertion and SV deletion within each continental group; error bars indicate one standard deviation. Colors denote continental groups. The pie chart shows the proportions of CpGs altered by each genetic variant class. **c**, Methylation levels of CpG gain and loss across genetic variant classes and continental groups. Points indicate the median methylation levels of CpGs associated with SNV gain, SNV loss, indel insertion, indel deletion, SV insertion and SV deletion within each continental group; error bars indicate one standard deviation. Colors denote continental groups. The dashed horizontal line indicates the genome-wide median methylation level across haplotype-resolved methylomes (80.3%). **d**, Density distributions of CpG methylation levels in the paternal and maternal haplotypes of HG002, stratified by genetic variant class and CpG gain or loss status. Curves show kernel density estimates of methylation levels for non-var-CpGs and CpGs associated with SNVs, indels or SVs. Non-var-CpGs were defined as CpGs that did not overlap genetic variants. Colors indicate CpG class: non-var-CpGs, black; SNV-associated CpGs, gray; indel-associated CpGs, brown; and SV-associated CpGs, teal. For var-CpGs, solid and dashed lines indicate CpG gains and losses, respectively. Vertical lines indicate the median methylation level of each distribution. **e**, Frequencies of CpG gain and loss events in HPRC2 samples. For SVs, each affected CpG was counted as a separate var-CpG event; therefore, a single SV could contribute multiple events. Pie charts show the proportions of var-CpGs with frequencies below 0.1, between 0.1 and 0.9, or above 0.9, stratified by variant class and CpG gain or loss status. Colors indicate frequency ranges: orange, <0.1; light blue, 0.1–0.9; and dark blue, >0.9. **f**, Transposable element (TE) composition of CpG gain and loss in SV-associated var-CpGs. Colors indicate TE categories: DNA transposons, LTRs, LINEs, SINEs, SVAs and Other; Other includes other TE classes and sites without TE annotation. **g**, Methylation levels of CpG gain and loss sites within TEs. Violin plots show median methylation levels of CpG sites gained by insertions and lost by deletions across transposable element classes. Each distribution represents haplotype-level median methylation values for CpG gain or loss sites within the indicated TE class. White diamonds indicate group medians. The dashed horizontal line indicates the genome-wide median methylation level across haplotype-resolved methylomes (80.3%). Paired comparisons between CpG gain and CpG loss were performed within each TE class using two-sided paired Wilcoxon signed-rank tests. ***, FDR < 0.001. **h**, Numbers of SV-driven CpG island (CGI) gains and losses per methylome. Points represent individual haplotype-resolved assemblies. **i**, Contribution of mobile element insertions (MEIs) to SV-driven CGI gains and losses. SV-driven CGI gain and loss events were categorized as SINE/AluY-driven CGIs, other MEI-driven CGIs and non-MEI-driven CGIs. **j**, HPRC epigenome browser view of the insertion on chromosome 2 in GRCh38 and the two haplotypes of HG02572, with tracks showing sequence alignment, RepeatMasker annotations, CGIs and PacBio methylation levels. HG02572 haplotype 2 (Hap2) carries an AluY insertion that is absent from GRCh38 and creates a hypermethylated CGI. **k**, Geographic distribution of allele frequencies for an AluY insertion across sampled continental groups. Circles indicate sampling locations and are colored according to allele frequency; continental group abbreviations are shown adjacent to each location. **l**, Population-associated expression of *NMI* and *RIF1* isoforms. Raincloud plots show laboratory-and sex-adjusted inverse-normalized transcripts per million (TPM) values for the *NMI* transcript flairiso38094-1 and the *RIF1* transcript ENST00000428287.6 in the AFR group and all other continental groups combined. Half-violin plots show expression distributions, points represent individual samples, and box plots indicate medians and interquartile ranges. Comparisons between the AFR group and all other continental groups were performed using Student’s t-tests.

SNVs accounted for most var-CpGs (∼76%), followed by SVs (∼22%) and indels (∼2%) (Fig. 2b). Although partly reflecting variant abundance, SVs were a substantial source of CpG gain and loss: approximately two-thirds of SVs contained at least one CpG (Extended Data Fig. 5a). SV insertions contributed 1.9-fold more var-CpGs than SV deletions, consistent with the higher CpG density of insertion SVs (Fig. 2b and Extended Data Fig. 5b).

Leveraging the panepigenome graph, we estimated the frequencies of var-CpG gain and loss events across HPRC2 haplotypes. The mean frequency was 4.09% overall, with SV-associated var-CpGs occurring at lower frequencies than SNV- and indel-associated var-CpGs (∼2% versus ∼5%), consistent with the generally lower frequencies of SVs^49^ (Fig. 2e). CpG gain events had lower frequencies than losses across variant classes, with the largest difference observed for SV-associated var-CpGs. Although most events were low-frequency, 141,815 occurred at frequency >90% relative to GRCh38 (Fig. 2e). Among these high-frequency events, 70.29% overlapped genes and 11.11% overlapped promoters, indicating that genetically driven CpG gain and loss frequently occur in regions with potential regulatory relevance^8,50,51^.

#### Structural variants and transposable elements shape var-CpG landscapes

SVs account for a large fraction of polymorphic bases in human genomes^49^ and contributed substantially to CpG gain and loss. Compared with SNVs and indels, SVs were 2.44- and 21.59-fold more likely to contain CpGs, respectively (Extended Data Fig. 5a). Although mostly reflecting their larger size, CpGs were also denser within SVs than in randomly sampled genomic regions (Extended Data Fig. 5b). SV-associated CpGs showed higher methylation than SNV- and indel-associated CpGs, by 13.8% and 10.1%, respectively, indicating a distinct methylation profile^25^ (Fig. 2c).

Because transposable elements (TEs) contribute substantially to SVs and are often highly methylated^52–54^, we examined TE-overlapping SV-associated CpGs. Overall, 47.7% of SV-associated CpG gains and 25.4% of losses overlapped major TE classes, with short interspersed nuclear elements (SINEs) accounting for approximately half of these events (Fig. 2f). TE-associated SV-CpGs were generally highly methylated, although levels varied across TE classes (Fig. 2g). Long terminal repeat (LTR)-associated losses approached the genome-wide median, whereas SINE-VNTR-Alu element (SVA)-associated gains showed particularly high methylation levels, potentially contributing to the elevated methylation of SV-associated CpGs overall.

CpGs associated with SV insertions showed ∼2-fold greater overlap with SVAs and SINEs than those associated with SV deletions (Fig. 2f). Because CpGs within these TE classes were among the most highly methylated, differences in TE composition may contribute to methylation differences between SV-associated CpG gains and losses (Fig. 2g).

#### Haplotype-resolved functional annotations of var-CpGs

To place var-CpGs in functional context, we annotated them using haplotype-resolved genomic features derived from HPRC2 assemblies^30^. Var-CpGs associated with different variant classes showed distinct genomic enrichments relative to all CpGs within each methylome (Extended Data Fig. 6a). SV-associated CpGs were more strongly enriched in regulatory features, including promoters and CGIs, than SNV- or indel-associated CpGs and were consistently more highly methylated (Extended Data Fig. 6b). For example, median methylation of SV-associated var-CpGs reached 87.8% in promoters and 92.4% in CGIs, compared with 25.7% and 44.4%, respectively, for indel-associated var-CpGs.

Approximately half of SV-associated CpG gains and losses occurred within CGIs (49.5–51.6%), representing ∼4.8-fold enrichment relative to all CpGs within each methylome (Extended Data Fig. 6a). Thus, rather than altering individual CpGs, SVs can create or remove entire CGIs. Each methylome contained an average of 784 SV- driven CGI gains and 757 losses (Fig. 2h and Supplementary Table 9). Across haplotypes, 83–91% of these CGI gains and losses were attributable to mobile element insertions (MEIs), with SINE/AluY elements accounting for >92% of MEI-driven events (Fig. 2i).

An AluY MEI created a CGI and was more frequent in AFR than in the other continental groups (minor allele frequency (MAF), ∼15% versus ∼2%; Fig. 2j,k). HPRC2 eQTL annotations linked this AluY insertion to expression of the immune-associated gene *NMI* and the DNA replication and repair factor *RIF1*; expression of both transcripts differed significantly between AFR and the other continental groups (Fig. 2l). This example illustrates how population-associated, SV-derived CGIs captured by haplotype-resolved genomes can reveal potential regulatory variation missed by conventional reference-based epigenomic analyses.

#### Haplotype-level population specificity of var-CpGs

Human demographic history is reflected in genome-wide differences in allele frequencies (AFs)^36,39–44^. Because var-CpG gain and loss are genetically determined, we examined whether var-CpG abundance and methylation also exhibited population-associated patterns across the five continental groups and 26 populations.

Consistent with global patterns of genetic diversity^36,41,44,49^, var-CpG counts differed significantly among continental groups and populations, with AFR methylomes containing the most var-CpGs, partly accounting for their greater number of non-reference CpGs (Fig. 2b). Population differences were substantially weaker for SV-associated CpGs than for SNV- or indel-associated CpGs, potentially reflecting the lower allele frequencies of SVs and stronger purifying selection against variants with larger functional effects^55^ (Fig. 2b).

In contrast to var-CpG abundance, which was highest in AFR haplotypes, var-CpG methylation was highest in AMR haplotypes and differed significantly across continental groups and populations (Kruskal–Wallis test, FDR < 0.05; Fig. 2c). As with abundance, continental differences in methylation were weaker for SV-associated than for SNV- or indel-associated var-CpGs (Supplementary Table 10). Together, these patterns indicate that var-CpG abundance and methylation capture complementary dimensions of population-associated epigenomic variation.

### Var-CpGs capture population-scale epigenomic diversity

Population-associated epigenomic diversity can arise from variation in both whether CpGs are present and their methylation levels when present.

#### Var-CpGs capture population-associated epigenomic diversity

To quantify epigenomic diversity beyond the linear reference genome, we defined three complementary layers: methylation at shared reference CpGs, CpG gain and loss at var-CpGs, and integrated var-CpG variation combining CpG presence or absence with methylation levels on CpG-containing haplotypes (Fig. 3a). Analyses included shared autosomal reference CpGs present in nearly all haplotypes and intermediate-frequency autosomal var-CpGs (5% < AF < 95%). The integrated var-CpG layer was modeled using a convolutional neural network (CNN) with an encoder–decoder architecture^56^ that jointly captured CpG presence or absence and, when present, its methylation level.

**Fig. 3:**
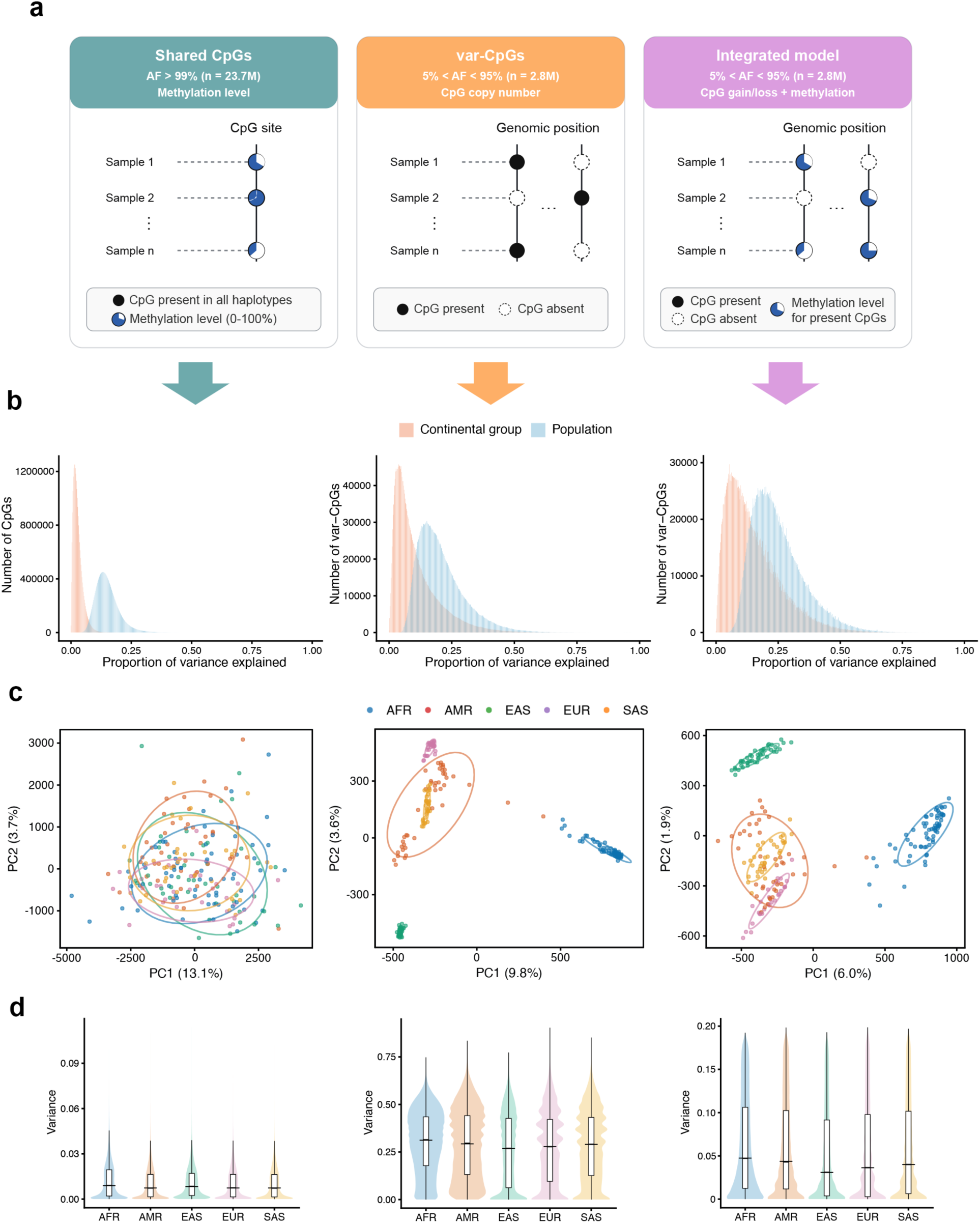
Patterns of panepigenomic diversity within and between populations. **a**, Three models quantify complementary layers of epigenomic diversity. From left to right, the models represent methylation variation at shared CpGs, CpG gain and loss at var-CpGs, and an integrated model combining CpG presence or absence with methylation variation at CpG-containing haplotypes. The integrated model was derived using a convolutional encoder–decoder architecture. **b**, Proportion of variance explained by continental group and population in each of the three panepigenomic models. From left to right, results are shown for methylation variation at shared CpGs, CpG gain and loss at var-CpGs, and the integrated model. **c**, Principal component analysis of the three panepigenomic models using the first two principal components (PCs). Points represent individuals and are colored by continental group. From left to right, results are shown for methylation variation at shared CpGs, CpG gain and loss at var-CpGs, and the integrated model. **d**, Variance in the three panepigenomic models across continental groups. From left to right, results are shown for methylation variation at shared CpGs, CpG gain and loss at var-CpGs, and the integrated model.

Following previous analyses of gene expression and splicing across human populations^57^, we fitted linear models to quantify variance explained by 1KG continental group and population labels^36^. At shared CpGs, continental group explained an average of 2.83% of methylation variance, whereas population explained 15.01% (Fig. 3b). Continental group effects did not exceed null expectations (one-tailed permutation test, *P* = 0.18), whereas population effects did (*P* < 0.001), indicating fine-scale population-associated methylation structure at shared CpGs.

Var-CpG-based models revealed pronounced population-associated structure. Continental group and population explained 15.10% and 24.97% of variance in CpG gain and loss, respectively, and 10.85% and 20.83% in the integrated model (Fig. 3b), with all effects exceeding null expectations (one-tailed permutation test, *P* < 0.001). Consistent with this, principal component analysis (PCA) of CpG gain and loss separated the five continental groups within the first three PCs, recapitulating broad genetic population structure^36,39–44^ (Fig. 3c and Extended Data Fig. 7). Although variance explained is not directly comparable across models because they quantify different biological features, PCA showed clearer continental structure for var-CpGs than for shared CpGs. CpG gain/loss and integrated models produced related but non-identical clustering, consistent with additional information from var-CpG methylation, although variant-conditioned analyses are required to quantify this contribution.

Because most human genetic variation occurs within rather than between populations^42^, we next examined inter-individual epigenomic variation within continental groups. Inter-individual variance differed significantly across continental groups in all three models (*P* < 2.2 × 10⁻¹⁶), with AFR showing the greatest diversity (Fig. 3d), consistent with greater genetic diversity in AFR and reduced diversity outside Africa following serial founder effects during human dispersal^57–59^. Together, var-CpGs capture population-associated epigenomic variation beyond that represented by shared reference CpGs alone.

#### Var-CpGs expand population-associated epigenomic signals

By representing CpG presence and absence across diverse haplotypes, the human panepigenome enabled detection of population-associated epigenomic variation beyond the linear reference genome. We analyzed shared CpGs and var-CpGs using pairwise continental group and focal-versus-rest comparisons. Unless otherwise indicated, population-specific signals were defined as |Δ| > 0.20 and FDR < 0.05, where Δ denotes the between-group difference in the mean of the corresponding measure. Only 0.02% of shared CpGs were population-specific per focal-versus-rest comparison, compared with 15.0% of var-CpGs in the CpG gain/loss model and 7.4% in the integrated model (Fig. 4a). Similar trends were observed using a nominal threshold of *P* < 5.0 × 10⁻⁸ and in pairwise continental group comparisons (Extended Data Fig. 8a,b).

**Fig. 4:**
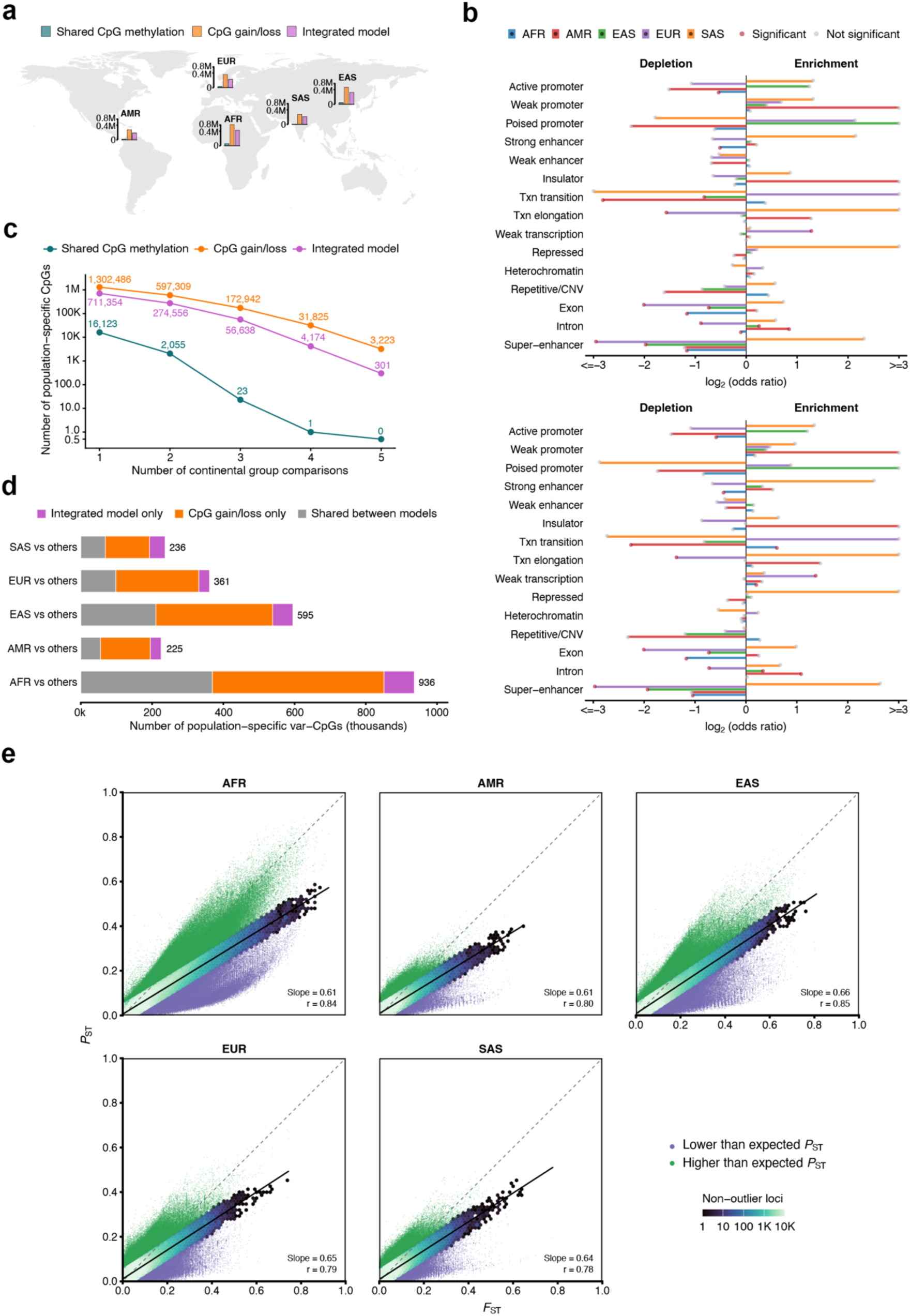
Panepigenomic analysis of human diversity. **a**, Global distribution of population-specific CpGs across continental groups. Bars positioned at schematic continental locations show the numbers of population-specific CpGs identified from methylation differences at shared CpGs, CpG gain or loss at var-CpGs, and the integrated model in focal-versus-rest comparisons for AFR, AMR, EAS, EUR and SAS. Bar heights are proportional to the square root of the corresponding CpG count, with local axes indicating the transformed scale. Colors denote analysis type: teal, methylation differences at shared CpGs; orange, CpG gain/loss at var-CpGs; and purple, integrated model. **b**, Genomic enrichment and depletion of population-specific var-CpGs relative to population-specific shared CpGs. The upper panel shows enrichment patterns for population-specific var-CpGs identified using the CpG gain/loss model, whereas the lower panel shows those identified using the integrated model. Population-specific CpGs were identified for each continental group in focal-versus-rest comparisons with |Δ| > 0.20 and nominal *P* < 5.0 × 10⁻⁸. Bars show the log₂ odds ratios for enrichment of population-specific var-CpGs in each genomic feature, separately for AFR, AMR, EAS, EUR and SAS compared with all other continental groups. Negative and positive values indicate depletion and enrichment, respectively. Bar colors denote continental groups. Red points indicate significant enrichment or depletion at FDR < 0.05, whereas gray points indicate nonsignificant enrichment or depletion. For visualization, log₂ odds ratios were capped at −3 and 3. Txn, transcription; CNV, copy number variation. **c**, Numbers of population-specific CpGs identified in one or more continental group comparisons for shared CpG methylation, CpG gain/loss and the integrated model. Population-specific CpGs were identified in focal-versus-rest comparisons with |Δ| > 0.20 and FDR < 0.05. **d**, Overlap of population-specific var-CpGs between the CpG gain/loss and integrated models. Population-specific CpGs were identified for each continental group in focal-versus-rest comparisons with |Δ| > 0.20 and FDR < 0.05. **e**, Relationship between genetic differentiation and integrated genetic–methylation differentiation of var-CpGs across continental groups. Hexagonal bins show the density of non-outlier var-CpGs according to genetic differentiation (*F*_ST_) and differentiation of the integrated genetic–methylation signal (*P*_ST_) in focal-versus-rest comparisons for AFR, AMR, EAS, EUR and SAS; darker colors indicate higher densities. Solid lines show linear regressions of *P*_ST_ on *F*_ST_ for each comparison and were used to estimate the expected *P*_ST_ at each locus; dashed diagonal lines indicate *P*_ST_ = *F*_ST_. Var-CpGs were classified as outliers when |observed *P*_ST_ − expected *P*_ST_| > 0.05. Loci with lower-and higher-than-expected *P*_ST_ are shown in purple and green, respectively. Outliers were excluded from the hexagonal-bin density layer and plotted separately as colored points.

Population-specific var-CpGs showed distinct genomic enrichments relative to population-specific shared CpGs across comparison designs and significance thresholds (Fig. 4b and Extended Data Fig. 9a–c). At |Δ| > 0.20 and *P* < 5.0 × 10⁻⁸, var-CpGs identified by the CpG gain/loss or integrated model were enriched in weakly transcribed regions in EUR-versus-rest comparisons and in introns in AMR-versus-rest comparisons (Fig. 4b). Enrichment patterns also differed between models: in AFR-versus-rest comparisons, enrichment in transcriptional transition and weakly transcribed regions was detected only by the integrated model. Population-specific var-CpGs were also more likely than population-specific shared CpGs to recur across multiple comparisons (39–46% versus 12.8% in focal-versus-rest analyses; Fig. 4c), with similar patterns across significance thresholds and pairwise comparisons (Extended Data Fig. 8c).

We refer to variants that create or disrupt CpGs as CpG-altering variants. Common CpG-altering variants (MAF > 1%) were 1.30-fold enriched at loci previously reported to be under directional selection^60^ relative to other common variants (Supplementary Table 11). These loci overlapped genes associated with human phenotypic diversity and population-differentiated disease risk, including pigmentation (*BNC2*, *KITLG*, *DSTYK*), hair morphology and male-pattern baldness (*TCHH*), vitamin D levels (*NADSYN1*), celiac disease (*TSBP1*, *HLA-DQB1*), Crohn’s disease (*SLC22A4*), gout (*ABCG2*) and hemochromatosis (*HFE*). Together, these findings link CpG-altering variation to loci associated with recent human adaptation and phenotypic diversity.

#### Genetic and epigenetic contributions to population-specific var-CpGs

We quantified the relative contributions of CpG gain/loss and methylation to population-associated var-CpG variation by comparing the CpG gain/loss and integrated models (Fig. 4d and Extended Data Fig. 8d,e). In focal-versus-rest comparisons, the CpG gain/loss model identified approximately twice as many population-associated var-CpGs as the integrated model, with ∼57% detected exclusively by CpG gain/loss (Fig. 4d). Thus, population-associated differentiation at var-CpGs was more often captured by CpG presence or absence alone than by the integrated genetic–methylation signal under these thresholds. Extending previous observations that population-associated methylation at reference CpGs is often associated with nearby genetic variants^15,17^, these findings show that genetic variation shapes the methylome not only by altering methylation at existing CpGs but also by creating or disrupting the CpG substrate itself.

Notably, ∼1.8% of population-associated var-CpGs were detected exclusively by the integrated model (Fig. 4d). One SV insertion-associated var-CpG within an intron of *ROBO1* was identified exclusively by this model in the AFR-versus-rest comparison and in all pairwise comparisons involving AFR. *ROBO1* has previously been implicated in tumor suppression and population-associated methylation differences in prostate cancer^61^. Thus, incorporating methylation reveals population-associated signals not captured by CpG presence or absence alone.

#### Genetic and epigenetic differentiation at var-CpGs

To quantify population-associated var-CpG divergence beyond binary identification of population-specific sites, we measured genetic differentiation using *F*_ST_ and integrated genetic–methylation differentiation using *P*ST, an *F*_ST_-analogous measure of quantitative differentiation^14,62^. Genome-wide, *F*_ST_ and *P*_ST_ were strongly correlated across var-CpGs in all focal-versus-rest comparisons (Pearson’s r = 0.78–0.85; Fig. 4e), indicating that integrated var-CpG differentiation broadly tracked underlying genetic differentiation. However, a subset deviated substantially from this relationship, with integrated differentiation either lower or higher than expected from the genome-wide relationship with genetic differentiation.

We estimated expected *P*_ST_ from the fitted linear regression of *P*_ST_ on *F*_ST_ within each comparison and classified loci as deviating from expectation when |observed *P*_ST_ − expected *P*ST| > 0.05. On average, ∼94,000 var-CpGs per comparison showed lower-than-expected *P*ST, consistent with attenuated integrated differentiation (Fig. 4e). A var-CpG associated with an SV insertion at *TRPV1* showed markedly lower *P*_ST_ than expected from its genetic differentiation in the AFR-versus-rest comparison (expected *P*_ST_ = 0.43; observed *P*_ST_ = 0.23). *TRPV1* encodes a cation channel involved in thermosensation, nociception and inflammation^63^. The AluY MEI described above, which created a CGI and was linked to *NMI* and *RIF1* transcript expression (Fig. 2j–l), contained multiple var-CpGs with similarly lower-than-expected *P*_ST_ in the AFR-versus-rest comparison; for example, one var-CpG had an expected *P*_ST_ of 0.16 and an observed *P*_ST_ of 0.05. Together, these examples illustrate methylation-associated attenuation of integrated differentiation at both an individual var-CpG and a CGI created by a single insertion, consistent with the hypothesis of epigenetic buffering relative to the underlying genetic variation^64^.

Conversely, ∼141,000 var-CpGs per comparison showed higher-than-expected *P*ST, indicating amplified integrated differentiation (Fig. 4e). An rs3184504-altered var-CpG in *SH2B3* showed higher-than-expected differentiation in EAS-versus-rest (expected *P*_ST_ = 0.07; observed *P*_ST_ = 0.15) and EUR-versus-rest comparisons (expected *P*_ST_ = 0.31; observed *P*_ST_ = 0.38). *SH2B3* is involved in immune signaling and immune-mediated disease, and the derived allele at rs3184504 has been reported to show signatures of recent positive selection^65^. Together, these findings indicate that var-CpG differentiation largely tracks underlying genetic variation, while methylation is associated with either attenuated or amplified differentiation at a subset of loci.

### Var-CpGs link sequence variation, methylation and transcription

The functional significance of var-CpGs depends on how CpG gain or loss and methylation connect genetic variation to gene regulation. We therefore examined associations among var-CpG methylation, local genetic variation and transcript expression.

#### Genetic associations with var-CpG methylation

Conventional methylation QTL (mQTL) analyses test genetic associations with methylation at existing CpGs^66,67^, whereas var-CpGs add another dimension because genetic variants can determine whether a CpG is present and influence its methylation level when present. We tested intermediate-frequency autosomal var-CpGs for methylation associations with (1) their own copy number, (2) nearby var-CpG copy number within 10 kb and (3) nearby non-CpG-altering variants within 10 kb, restricting analyses to individuals carrying at least one copy of the CpG (Fig. 5a).

**Fig. 5:**
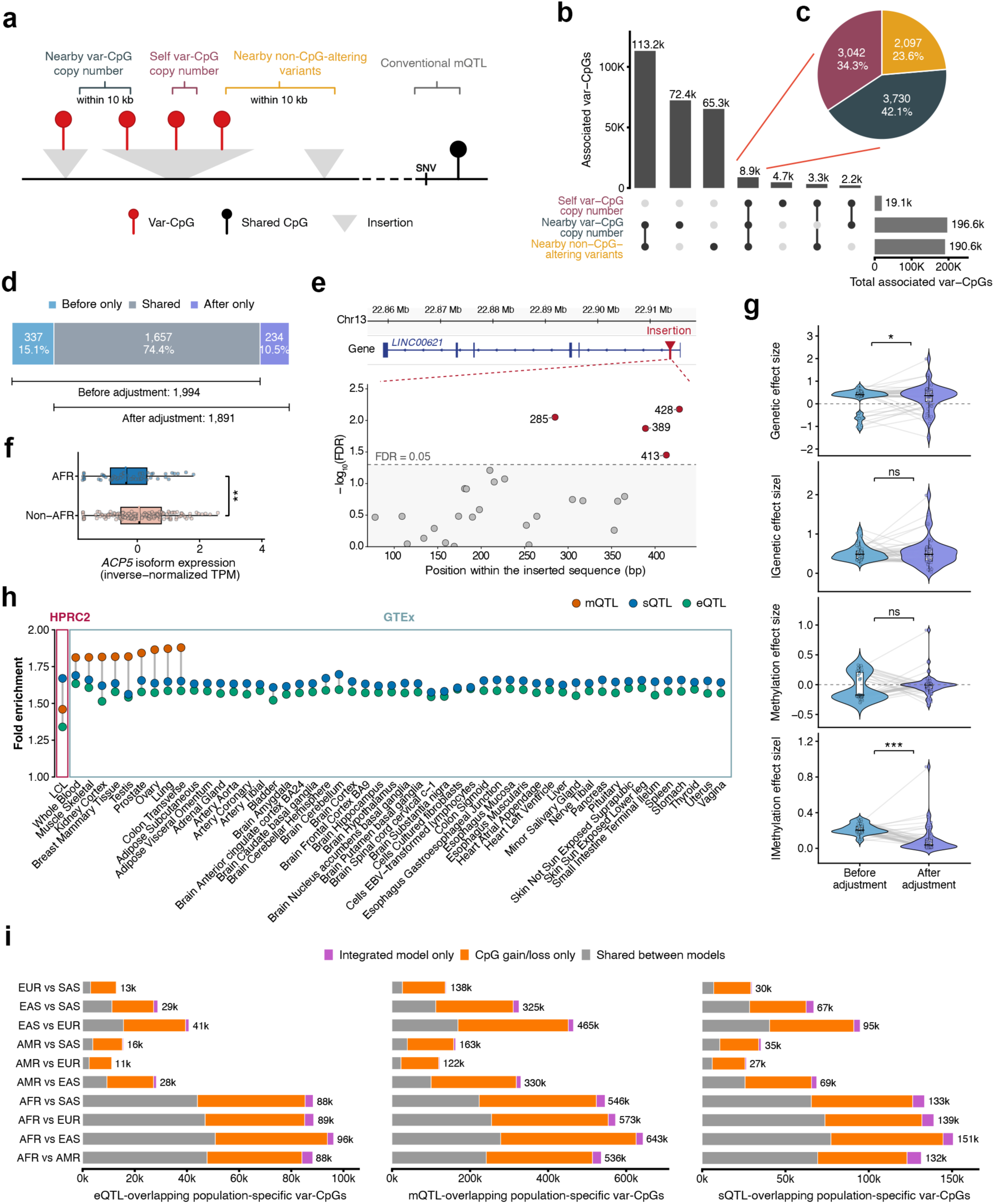
Var-CpGs link genetic variation, methylation and transcription. **a**, Schematic illustration of potential genetic effects on var-CpG methylation. Three classes of genetic features were tested: copy number of the var-CpG itself, copy number of nearby var-CpGs (within 10 kb), and nearby genetic variants (within 10 kb^17,18,96^) that do not alter CpG gain or loss. **b**, UpSet plot showing the number and overlap of var-CpGs whose methylation was associated with each of the three classes of genetic features. **c**, Pie chart showing relative contributions of the three classes of genetic features to shared var-CpG methylation. For each of the 8,869 var-CpGs shared across all three classes of genetic features, the genetic feature with the largest absolute regression coefficient (|β|) was identified. Percentages indicate the proportion of shared var-CpGs for which each genetic feature class exhibited the largest absolute effect size. **d**, Overlap of significant var-CpG–transcript pairs before and after adjustment for the underlying CpG-altering variant. The horizontal bar shows pairs detected only before adjustment, shared between analyses and detected only after adjustment. Percentages are calculated relative to the union of pairs identified in either analysis. Brackets indicate the total numbers detected before and after adjustment. **e**, Insertion-associated var-CpG signals at the *LINC00621* locus. Top, GRCh38 gene structure and insertion breakpoint at chr13:22,913,826. Bottom, association significance between var-CpGs along the inserted sequence and *LINC00621* transcript expression. Filled circles represent individual var-CpGs; red and gray indicate significant and nonsignificant associations, respectively. The dashed line marks FDR = 0.05, and labels indicate significant var-CpG positions. Upper and lower coordinates denote GRCh38 genomic positions and positions within the inserted sequence, respectively. **f**, Box plot showing population-associated *ACP5* transcript expression differences for the mediation-associated var-CpG. Laboratory- and sex-adjusted inverse-normalized transcripts per million (TPM) values for the *ACP5* transcript ENST00000695812.1 are shown for AFR and non-AFR individuals at a promoter var-CpG altered by rs111931834. *P* values were calculated using Student’s t-test. **, *P* < 0.01. **g**, Box plots show genetic and methylation effect sizes before and after adjustment for variant-by-methylation interaction among pairs with significant interaction effects. Signed β values are shown in the top panels and absolute β values in the bottom panels. Lines connect the same var-CpG–transcript pair across models. *P* values were calculated using two-sided paired Wilcoxon signed-rank tests. *, *P* < 0.05; ***, *P* < 0.001; ns, nonsignificant (*P* ≥ 0.05). **h**, CpG-altering variants are enriched in HPRC2 and GTEx molecular QTLs. Fold enrichment was calculated by comparing CpG-altering variants with other variants, and significance was assessed using Pearson’s χ² test followed by Benjamini–Hochberg correction. Significant enrichment was defined as FDR < 0.05. eQTL, expression quantitative trait locus; mQTL, methylation quantitative trait locus; sQTL, splicing quantitative trait locus. **i**, Population specificity of CpG-altering variants overlapping HPRC2 molecular QTLs. Numbers of CpG-altering variants overlapping HPRC2 eQTLs, mQTLs and sQTLs that were classified as population-specific by either the CpG gain/loss or integrated model in each pairwise comparison between continental groups. Population-specific var-CpGs were defined using ∣Δ∣ > 0.20 and FDR < 0.05.

Overall, methylation levels of ∼270,000 var-CpGs (9.58%) were associated with at least one genetic feature class, most frequently with nearby var-CpGs or non-CpG-altering variants (Fig. 5b). Of these, 8,869 were associated with all three classes, whereas others showed class-specific associations. These findings reveal distinct genetic associations involving the underlying CpG-altering variant, neighboring var-CpGs and nearby non-CpG-altering variants, broadening the genetic contexts considered beyond conventional mQTL analyses^66,67^.

#### Coupling between CpG copy number and methylation at regulatory loci

Among var-CpGs whose methylation was significantly associated with all three genetic feature classes, nearby var-CpG copy number most frequently showed the largest absolute regression coefficient (|β|), followed by self-copy number and nearby non-CpG-altering variants (Fig. 5c). Thus, CpG copy number features generally showed larger methylation associations than nearby variants that did not alter CpG content.

The direction of association further supported coupling between CpG copy number and methylation. Positive associations predominated for self-copy number and nearby var-CpG copy number, particularly for var-CpGs associated with all three genetic feature classes (99.55% and 62.61%, respectively; Extended Data Fig. 10a).

Compared with var-CpGs associated with at least one genetic feature class, those associated with all three classes were enriched in transcriptionally active chromatin, enhancers and genic regions, and depleted in heterochromatin, repressed chromatin and repetitive or copy-number-variable regions (FDR < 0.05; Extended Data Fig. 10b). These multilayer genetic associations therefore occurred preferentially within active regulatory contexts. Together with evidence that CpG deamination contributes to enhancer evolution^68,69^, these findings suggest that genetic variation affecting CpG content may contribute to regulatory evolution through linked variation in CpG dosage and methylation at genetically variable loci.

#### Promoter var-CpG methylation is associated with transcript expression

Expression quantitative trait methylation (eQTM) analyses test associations between DNAm and gene or transcript expression^70–72^. Although promoter methylation at reference CpGs is often associated with transcriptional repression^8,50,51^, whether promoter var-CpG methylation is associated with transcript expression remains unclear. Across 195 individuals with matched methylomes and Iso-Seq data, we identified 1,994 significant var-CpG–transcript pairs involving 960 promoter var-CpGs and 804 transcripts (Fig. 5d). Most associations were negative (64.5%), consistent with canonical promoter methylation–transcription relationships^73,74^. Distinct var-CpGs within the same SV showed different associations with the same transcript, revealing regulatory variation at single-CpG resolution (Fig. 5e and Supplementary Table 12).

After adjusting for the underlying CpG-altering variant, 1,891 var-CpG–transcript pairs were significant (Fig. 5d). Of these, 1,657 (87.6%) were also significant before adjustment, indicating that most associations persisted after accounting for the underlying variant, whereas a smaller subset emerged only after adjustment. Thus, most promoter var-CpG–transcript associations could not be attributed solely to the underlying CpG-altering variant alone, extending eQTM analysis to genetically variable CpG substrates.

#### Mediation analysis identifies potential regulatory relationships at selected var-CpGs

We next tested whether var-CpG methylation statistically mediated associations between underlying CpG-altering variants and transcript expression. Mediation analysis identified 13 significant events involving nine var-CpGs and nine transcripts (Supplementary Table 13). A promoter var-CpG in *ACP5*, altered by rs111931834, statistically mediated the association between the variant and ENST00000695812.1 expression. Rs111931834 has been associated with circulating tartrate-resistant acid phosphatase type 5 levels, whereas *ACP5* is implicated in spondyloenchondrodysplasia with immune dysregulation^75,76^. Given the limited sample size and small number of significant events, these findings suggest that var-CpG methylation may statistically mediate genetic effects on transcript expression at selected loci.

Eight of nine mediation-associated var-CpGs showed population-associated patterns (Supplementary Table 14). The *ACP5* var-CpG was less frequent in AFR than in EAS (Δ = −0.22), accompanied by modest differences in methylation and transcript expression (Fig. 5f). Together, these findings identify a subset of var-CpGs at which methylation statistically mediates associations between genetic variation and transcript expression, with most also showing population-associated variation.

#### Variant-by-methylation interactions at var-CpGs in gene regulation

Whereas mediation tests whether methylation statistically mediates genetic effects on transcription, interaction analysis tests whether the association between methylation and transcription depends on the underlying CpG-altering variant state. Previous studies have shown that methylation at reference CpGs can interact with nearby genetic variation to influence gene expression^77^. We therefore tested promoter var-CpGs for variant-by-methylation interactions influencing transcript expression. We identified 46 significant var-CpG–transcript pairs involving 25 var-CpGs (0.03% of those tested) and 32 transcripts (Supplementary Table 15).

Among significant pairs, absolute methylation effect sizes were reduced after including the interaction term, whereas signed methylation effects did not differ significantly, indicating that some methylation associations were variant-dependent (Fig. 5g). Consistently, methylation effects at most loci were no longer significant after accounting for the interaction. One exception was an rs1764287-generated promoter var-CpG in *ZNHIT6*, where methylation was positively associated with ENST00000370574.4 expression (β = 0.91), whereas an opposing interaction effect (β = −0.38) attenuated this association.

We next assessed whether variant-by-methylation interactions altered the genetic effect on transcript expression. Absolute genetic effect sizes did not differ significantly between models with and without the interaction term, whereas signed genetic effects did, consistent with locus-specific changes in effect direction or magnitude (Fig. 5g). Among significant transcript–var-CpG pairs, interaction terms attenuated genetic effects in 60.87% of cases, although reinforcing effects were also observed (Supplementary Table 15). For example, at an indel-associated var-CpG in *CYP4V2*, the genetic term was positively associated with expression of transcript flairiso123638-4 (β = 0.61), whereas the interaction coefficient was negative (β = −0.17), consistent with attenuation of the genetic effect and further supporting the epigenetic buffering hypothesis. Together, these results identify locus-specific variant-by-methylation interactions in transcript regulation.

#### CpG-altering variants are enriched in molecular QTLs

Having linked promoter var-CpG methylation to transcript expression at specific loci, we next asked whether CpG-altering variants were broadly enriched among molecular QTLs (xQTLs). Among common variants (MAF > 1%), CpG-altering variants were significantly enriched among eQTLs, mQTLs and sQTLs identified in HPRC2 LCLs^78^ (Fig. 5h). Similar enrichments were observed across GTEx tissues^67,79^, ranging from 1.51–1.63-fold for eQTLs, 1.81–1.88-fold for mQTLs and 1.56–1.70-fold for sQTLs. Thus, gain or loss of CpG sites may contribute to molecular QTL architecture by altering the substrate available for DNAm.

Because molecular QTL effects can vary across populations^80^, we examined population-associated variation among HPRC2 xQTL-associated var-CpGs. More than three-quarters of eQTL-, sQTL- and mQTL-associated var-CpGs were population-associated in at least one pairwise continental group comparison (77.5–81.3%). Although the CpG gain/loss model identified most signals across xQTL classes, the integrated model exclusively detected an additional 9.7–13.4% (Fig. 5i).

Among signals unique to the integrated model, 4,392 overlapped all three xQTL classes, including 73 at loci previously implicated in directional selection^60^ (Supplementary Table 16). An rs6568949-altered var-CpG showed an AMR–EAS-specific signal detected only by the integrated model. rs6568949 is an eQTL for *CALHM6*, which is involved in the early innate immune response to *Listeria monocytogenes* infection^81^. Together, these findings show that CpG-altering variants are enriched at molecular regulatory loci and that incorporating var-CpG methylation reveals additional population-associated regulatory signals.

### Clinical and pharmacogenomic relevance of var-CpGs

If CpG-altering variants contribute to regulatory variation, they may also intersect loci associated with human disease and therapeutic response.

#### CpG-altering variants are enriched for clinical and pharmacogenomic annotations

Having linked var-CpGs to transcript regulation and molecular QTLs, we next examined clinically and pharmacogenomically annotated variants. Among common CpG-altering variants in HPRC2 (MAF > 1%), 48,892 (1.08%) were annotated in ClinVar^82^, representing a 1.79-fold enrichment relative to other common variants (Fig. 6a). Enrichment varied between CpG gain and loss variants and across ClinVar categories, with the strongest enrichment among risk variants. Consistent with the largely healthy 1KG cohort^36^, most annotated variants were benign/likely benign, with only 7 pathogenic/likely pathogenic and 50 risk variants (Supplementary Table 17). We also identified 549 CpG-altering variants annotated in ClinPGx^83^, representing a 1.74-fold enrichment relative to other common variants (Fig. 6a and Supplementary Table 18). Together, these findings show that CpG-altering variants are disproportionately represented among clinically and pharmacogenomically annotated variants.

**Fig. 6:**
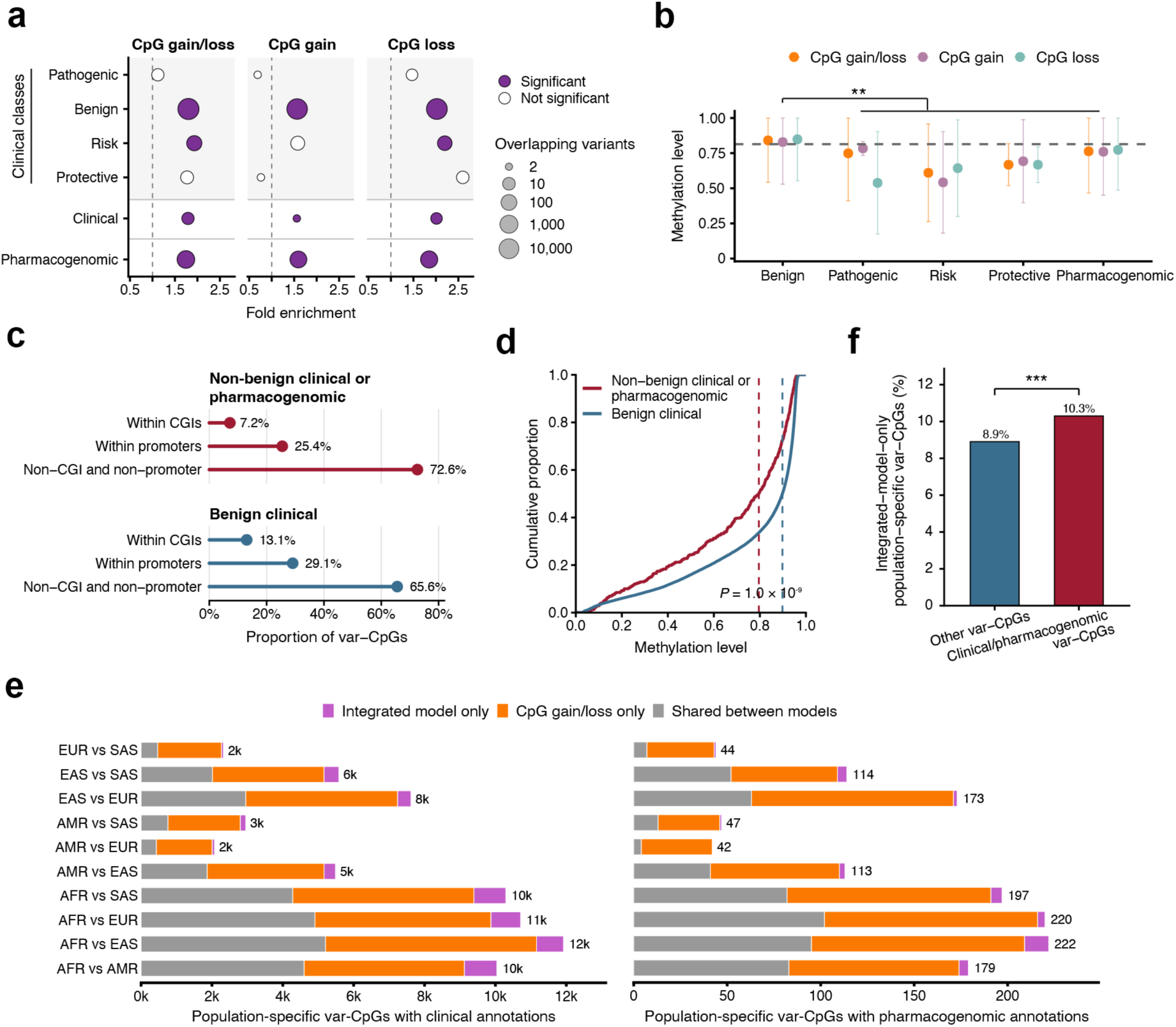
Clinical and pharmacogenomic relevance of var-CpGs. **a**, Enrichment of CpG-altering variants across clinical and pharmacogenomic annotation categories. Fold enrichment of CpG-altering variants among pathogenic, benign, risk, protective, overall clinical and pharmacogenomic variant annotations. Results are shown separately for all CpG-altering variants, CpG-gain variants and CpG-loss variants. Point size represents the number of overlapping variants. Purple points indicate significant enrichment or depletion at FDR < 0.05, whereas open points indicate nonsignificant associations. The dashed vertical line denotes no enrichment (fold enrichment = 1). Significance was assessed using Fisher’s exact test followed by Benjamini–Hochberg correction. **b**, Methylation patterns of clinically and pharmacogenomically annotated CpG-altering variants. Points indicate median methylation levels, stratified by annotation category and CpG gain or loss status; error bars indicate s.d. The dashed line indicates the genome-wide median methylation level of common var-CpGs (81.4%). Differences between groups were assessed using a two-sided Wilcoxon rank-sum test. **, *P* < 0.01. **c**, Genomic distribution of benign clinical var-CpGs and combined non-benign clinical and pharmacogenomic var-CpGs. Proportions of var-CpGs altered by benign clinical variants or by non-benign variants, including pathogenic, risk and protective variants, together with pharmacogenomic variants, that overlapped CpG islands (CGIs), promoters or regions outside both annotations. Points indicate the observed proportions, horizontal lines extend from zero to each estimate and labels show the corresponding percentages of var-CpGs. CGI and promoter annotations were analyzed independently and may overlap. **d**, Methylation distributions of benign clinical var-CpGs and combined non-benign clinical and pharmacogenomic var-CpGs. Empirical cumulative distribution functions show methylation levels for var-CpGs altered by benign clinical variants and for var-CpGs altered by non-benign variants, including pathogenic, risk and protective variants, together with pharmacogenomic variants. Dashed vertical lines indicate group medians. Differences between groups were assessed using a two-sided Wilcoxon rank-sum test. **e**, Population-specific clinically and pharmacogenomically annotated var-CpGs across continental groups. Population-specific var-CpGs were defined as sites with |Δ| > 0.20 and FDR < 0.05 in either the CpG gain/loss or integrated model in pairwise comparisons between continental groups. **f**, Enrichment of integrated-model-only population-specific var-CpGs among clinically or pharmacogenomically annotated var-CpGs. Bars show the proportions of integrated-model-only population-specific var-CpGs among clinically or pharmacogenomically annotated var-CpGs and among all other var-CpGs. Labels indicate the corresponding percentages in each group. Fold enrichment was calculated as the ratio of the two proportions, and significance was assessed using Fisher’s exact test. ***, *P* < 0.001.

#### Clinical and pharmacogenomic var-CpGs show distinct methylation patterns

Var-CpGs altered by these annotated variants showed distinct methylation profiles. Non-benign clinical (pathogenic/likely pathogenic, risk and protective) and pharmacogenomic var-CpGs were generally less methylated than benign/likely benign var-CpGs (61–76% versus 84%; Fig. 6b). Because CGIs and promoters are typically CpG-rich and hypomethylated^84^, we tested whether genomic context explained these differences. Var-CpGs altered by benign variants were 1.82-fold enriched in CGIs relative to those altered by non-benign clinical or pharmacogenomic variants, whereas promoter enrichment did not differ significantly (Fig. 6c). After excluding promoter- and CGI-overlapping sites, non-benign clinical and pharmacogenomic var-CpGs remained significantly less methylated than benign/likely benign var-CpGs (Fig. 6d), indicating that their reduced methylation was not explained solely by genomic distribution. The lower methylation of non-benign var-CpGs may reflect differences in their genetic and regulatory contexts, consistent with patterns reported for trait-associated mQTLs^67^.

Variant-dependent methylation may further modulate the regulatory effects of CpG-altering variants. For example, the benign variant rs7576384 generated a var-CpG whose variant-by-methylation interaction attenuated its genetic effect on the *PAX8-AS1* transcript ENST00000613966.1 (variant β = 0.83; interaction β = −0.15), consistent with a potential epigenetic buffering effect on variant-associated regulation.

#### Population specificity of clinical and pharmacogenomic var-CpGs

We next examined population-specific patterns among clinically and pharmacogenomically annotated var-CpGs. Population-specific signals were observed for 64.2% of clinically annotated and 72.7% of pharmacogenomic var-CpGs, with most detected by the CpG gain/loss model and a subset by the integrated model (Fig. 6e). Notably, 10.35% and 7.02%, respectively, were detected exclusively by the integrated model; these annotated var-CpGs were 1.16-fold more likely than other var-CpGs to show integrated-model-exclusive signals (Fig. 6f and Supplementary Table 19). One example was an rs3792876-altered var-CpG in *SLC22A4*, detected exclusively by the integrated model in AFR–EAS and AMR–EAS comparisons. rs3792876 is a ClinVar risk variant associated with rheumatoid arthritis. Thus, integrating CpG gain/loss with methylation adds population-associated epigenomic context to clinically and pharmacogenomically annotated variation.

## Discussion

The human methylome cannot be defined solely by methylation levels at reference CpGs, because genetic variation also determines whether CpG substrates exist. By anchoring CpG presence or absence and methylation level to single-nucleotide graph coordinates across diverse haplotypes, the panepigenome resolves CpGs in non-reference and structurally variable regions that are absent from or incompletely represented in GRCh38. It therefore captures a layer of human epigenomic diversity largely missed by single haplotype, linear reference methylome maps.

This draft panepigenome integrates 440 long-read methylomes, haplotype-resolved assemblies and graph coordinates across 26 populations, revealing 12.8 million CpGs absent from GRCh38 and resolving their presence, absence and methylation across diverse haplotypes. Although currently limited to 5mC in LCLs and the diversity represented by HPRC2, the framework can be extend to other tissues and cell types, developmental stages, epigenetic marks and future pangenome releases.

Beyond its value as a resource, the panepigenome captures three complementary layers of epigenomic variation: methylation variation at shared CpGs, CpG gain or loss at var-CpGs, and an integrated representation combining CpG presence or absence with methylation. Integrated var-CpG differentiation broadly tracks underlying genetic differentiation but is attenuated or amplified at a subset of loci. Thus, population-associated epigenomic variation largely mirrors genetic structure, while methylation is associated with additional differentiation at selected loci. Enrichment of CpG-altering variants at loci with signatures of directional selection further links genetically driven CpG variation to population-associated regulatory diversity.

SVs are an important source of genetically driven CpG gain and loss. Rather than altering individual sites, SVs can create or remove entire CGIs, frequently through AluY MEIs. These events produce coordinated changes in CpG content and methylation across CpG-rich elements, extending the functional interpretation of SVs beyond effects on sequence dosage, gene structure and regulatory-element positioning. As long-read sequencing reveals further SV diversity^41,48,49^, integrating methylation into SV annotation may help prioritize structurally variable regions with potential regulatory effects.

Var-CpGs also link sequence diversity, methylation variation and transcript regulation. Extending mQTL and eQTM analyses to genetically variable CpG substrates enables CpG gain or loss, methylation and transcript expression to be analyzed jointly. A longstanding challenge in eQTM interpretation is distinguishing methylation-associated effects from those arising because the same or linked genetic variants influence both methylation and expression^15,77^. Most promoter var-CpG–transcript associations persisted after adjustment for the underlying CpG-altering variant, suggesting regulatory signals beyond the underlying sequence change. Smaller subsets showed statistical mediation or variant-by-methylation interactions, revealing locus-specific regulatory configurations that require functional validation.

Epigenetic buffering refers to the maintenance of epigenomic or regulatory states despite underlying sequence variation, thereby attenuating its molecular consequences^64^. Our analyses provide population-scale evidence consistent with this model: many var-CpGs showed lower integrated differentiation than expected from their genetic differentiation, and variant-by-methylation interactions more often attenuated than reinforced the genetic effects on transcript expression. This interpretation does not imply that methylation alters the genetic variant itself or establishes reverse causality; rather, it indicates that epigenomic and regulatory differentiation need not scale proportionally with genetic differentiation. Together, these findings suggest that DNAm may contribute to regulatory robustness at genetically variable loci, although targeted perturbation will be required to establish a causal buffering mechanism.

CpG-altering variants are enriched among eQTLs, mQTLs and sQTLs in HPRC2 LCLs and across GTEx tissues (1.34–1.88-fold), suggesting that gain or loss of the CpG substrate may contribute to molecular QTL architecture. Integrating genetic variation with epigenetic state may therefore provide a more complete view of regulatory variation across tissues and populations. CpG-altering variants were also enriched among clinically and pharmacogenomically annotated loci, and the integrated model identified population-associated signals at a subset not detected by CpG gain or loss alone. Incorporating var-CpGs into variant interpretation may therefore provide additional regulatory and population context, particularly in non-reference and structurally variable regions. These associations remain hypothesis-generating and do not establish disease mechanisms or clinical utility.

Several limitations should be considered. LCLs enable standardized population-scale comparisons but do not fully represent primary tissues or dynamic developmental and environmental contexts^14,17,85^. Continental groupings simplify continuous and admixed patterns of human ancestry, and some findings may depend on cohort composition. Mediation and interaction analyses provide statistical rather than experimental evidence of regulatory mechanisms. Extending this framework to primary tissues, larger and more diverse cohorts, disease-relevant systems, developmental stages and other epigenetic marks, together with functional perturbation, will help establish the generality and functional consequences of CpG gain, loss and var-CpG methylation.

More broadly, this work extends pangenomic representation from sequence variation toward regulatory variation. Although demonstrated here for DNAm, the underlying principle may extend across epigenomic analyses: genetic variation can alter not only the level or state of an epigenetic mark but also whether its genomic substrate exists and in what sequence context. By jointly representing CpG gain, loss and methylation, the panepigenome places genetic and epigenetic variation within a common framework. As pangenome resources expand, integrating epigenomic information should clarify how genetic diversity shapes regulatory diversity. Together, this work establishes a foundation for moving beyond single-reference methylome maps toward population-scale, haplotype-resolved panepigenomic maps that connect sequence diversity with epigenomic and regulatory variation beyond the linear reference genome.

## Methods

### HPRC sequencing and genome assembly

We analyzed lymphoblastoid cell line (LCL) samples from 231 individuals included in the Human Pangenome Reference Consortium release 2 (HPRC2), representing 26 populations across five continental groups^30^ : Africa (AFR, *n* = 70), the Americas (AMR, *n* = 44), East Asia (EAS, *n* = 51), Europe (EUR, *n* = 30) and South Asia (SAS, *n* = 36). Continental group and population labels were assigned according to the 1000 Genomes Project (1KG)^36^.

Two haplotype-resolved assemblies per individual (462 assemblies from 231 individuals) were obtained from HPRC2 and were generated from PacBio HiFi and Oxford Nanopore Technologies ultra-long reads using trio- or Hi-C-based phasing^30^.

### CpG site and CGI annotation

CpG sites were identified by locating all occurrences of the dinucleotide CG in each haplotype-resolved assembly, treating uppercase and lowercase sequences equivalently. CpG density was calculated as the number of CpG dinucleotides per kilobase of non-N sequence. Non-reference CpGs were defined as CpGs present in a haplotype-resolved assembly but absent from the GRCh38 backbone after projection to HPRC2 graph coordinates.

CpG islands (CGIs) were annotated separately in each haplotype assembly using the UCSC cpg_lh executable, downloaded from the UCSC software repository (http://hgdownload.soe.ucsc.edu/admin/exe/linux.x86_64/cpg_lh). The algorithm identifies maximally scoring segments by assigning a score of +17 to each CG dinucleotide and −1 to all other nucleotides. Candidate regions were retained if they had GC content ≥50%, length >200 bp and an observed-to-expected CpG ratio >0.6, following the criteria of Gardiner-Garden and Frommer^86^. CGI detection was performed across the complete assembly sequence without excluding soft-masked regions. CGI shores were defined as the 2-kb intervals immediately flanking each CGI, and CGI shelves as the subsequent 2-kb intervals. CGIs in GRCh38 and T2T-CHM13 were annotated using the same pipeline.

### Quality control of HiFi cytosine methylation data

We performed quality control (QC) on 1,154 HiFi BAM files from 231 individuals. The QC workflow comprised three stages: detection of technical outliers, assessment of batch effects and verification of genomic coverage (Extended Data Fig. 2).

First, technical outliers were identified from the median and distribution of ML-tag values in the raw reads. Reads were aligned to the T2T-CHM13 v2.0 reference genome (GCF_009914755.1) using pbmm2 v1.14.99 with --preset HIFI --unmapped. BAM files were retained if more than 80% of reads had a mapping quality (MAPQ) greater than 30, more than 90% had a CCS read-quality score greater than 0.99, the mean alignment identity exceeded 99%, the mean read length exceeded 10 kb and the overall mapping rate exceeded 99%. Median methylation values and methylation distribution profiles were additionally assessed to identify potential technical biases, yielding 1,107 BAM files from 226 individuals that passed file-level QC.

To assess batch effects, 5mC levels were quantified at single-base resolution using pb-CpG-tools with --min-mapq 10 and --pileup-mode model. Potential effects of sequencing platform (Sequel II versus Revio), sequencing time point and SMRT cell position were evaluated using technical replicates. Technical reproducibility was assessed separately using technical replicates from our long-read methylation dataset and from the previously published WGBS dataset^26^. Within each dataset, differentially methylated regions (DMRs) and differentially methylated loci (DMLs) between technical replicates were identified using the DSS R package^87^. Three BAM files from HG002 showing significant laboratory-associated differences were excluded (Wilcoxon rank-sum test, *P* = 0.008). The complete long-read methylation QC workflow is available on GitHub (https://github.com/functionalepigenomics/longread-methylation-qc).

Finally, the remaining 1,104 BAM files were merged by individual, and genomic coverage was assessed across 226 individuals. Six individuals with genomic coverage of 30× or less were excluded, yielding a final dataset of 1,092 BAM files from 220 individuals (AFR, *n* = 66; AMR, *n* = 43; EAS, *n* = 46; EUR, *n* = 30; SAS, *n* = 35).

### Phasing, alignment and methylation profiling

PacBio HiFi reads were aligned independently to the two haplotype-resolved assemblies for each individual using minimap2 (ref.^88^). For trio-phased assemblies, these haplotypes corresponded to paternal and maternal assemblies; otherwise, they were designated haplotypes 1 and 2. Reads were assigned to haplotypes by comparing alignment support between the two assemblies. For each read alignment, we extracted the read identifier, mapping quality, alignment score (mg) and the CCS read quality tag (rq). Reads were retained if rq ≥ 0.9 and mapping quality ≥ 10. For reads with multiple alignments within the same haplotype assembly, the best-supported alignment was retained. Alignment support was calculated as the product of the mg value and a read length consistency score, defined as the aligned read length divided by the total length of that read. Reads aligning to both haplotypes were assigned to the haplotype with stronger alignment support, whereas reads with equivalent support were randomly assigned between haplotypes to maintain balanced coverage for downstream analyses.

Haplotype-assigned BAM files were sorted using SAMtools v1.21. 5mC levels were quantified at single-base resolution using pb-CpG-tools v2.3.2 with --min-mapq 10 and --pileup-mode model, which estimates methylation levels from the MM and ML tags using a machine-learning model. To minimize bias from mismatches between reads and haplotype assemblies, we retained only CpGs present in the corresponding haplotype assembly and supported by a minimum coverage depth of 10×.

### Haplotype-resolved genomic feature and repeat annotations

RepeatMasker, centromeric satellite (CenSat), segmental duplication, transposable element (TE) and gene annotations for the HPRC2 assemblies were obtained from https://github.com/human-pangenomics/hprc_intermediate_assembly/tree/main/data_tables/annotation and were generated as described in the HPRC2 study^30^. Promoters were defined separately for each haplotype-resolved assembly using haplotype-specific gene annotations. Each promoter was defined as the strand-aware 2-kb interval upstream of the transcription start site (TSS) and clipped to contig boundaries. BEDTools v2.29.1 (ref.^89^) was used to intersect these genomic annotations with non-reference CpGs and variant-associated CpGs (var-CpGs).

### Projection of individual assembly coordinates to graph coordinates

Individual assembly coordinates were projected to HPRC2 graph coordinates using haplotype-specific walks encoded in the GFA representation of the Minigraph-Cactus graph, constructed with GRCh38 as the backbone. For each chromosome, the corresponding haplotype walk was parsed into an ordered series of oriented graph segments, and cumulative segment offsets were calculated along the walk. Input intervals, including CpG intervals, were sorted by assembly coordinate and located within this cumulative coordinate system. Each interval was represented by the traversed graph path, the start offset within the first segment, the end offset within the last segment and the total interval length. For intervals spanning multiple graph segments, the complete intervening path was retained. Segment orientation was accounted for explicitly, and sequences traversing reverse-oriented segments were reverse-complemented during reconstruction of interval sequences. Projected interval lengths and sequences were compared with those of the corresponding intervals in the source assemblies to verify coordinate consistency.

### CpG presence frequency estimation

For each graph CpG coordinate, presence frequency was calculated as the number of haplotypes carrying the CpG divided by the 440 haplotypes analyzed. CpGs were classified as rare (<1%), low frequency (1% to <10%), intermediate frequency (10% to <90%) or high frequency (≥90%). CpGs mapped to the GRCh38 backbone were classified as reference CpGs, whereas those not mapped to the backbone were classified as non-reference CpGs.

### Identification and classification of var-CpGs

Haplotype- and graph-based genetic variants, including SNVs, indels (1–50 bp) and structural variants (SVs; >50 bp), were obtained from the HPRC2 variant releases and were generated, integrated and filtered as described in the HPRC2 paper^30^. Haplotype-based variants were represented relative to GRCh38, whereas graph-based variants were obtained from the HPRC2 Minigraph-Cactus graph constructed with GRCh38 as the backbone. Graph variants were preprocessed using a modified decomposition workflow originally developed for PanGenie^90^.

var-CpGs were identified by intersecting CpG coordinates with variant-altered sequence intervals, including variant junctions. Haplotype-based intersections were performed using BEDTools v2.29.1 (ref.^89^), and graph-based intersections were performed as described below under “Graph-based interval-overlap analysis.” CpG gain and loss were determined from changes in the local dinucleotide context associated with each variant. A CpG gain was defined when the variant-containing haplotype carried a CG dinucleotide absent from GRCh38, including CpGs located within an inserted sequence or spanning a junction between variant and flanking sequences. A CpG loss was defined when a CG dinucleotide present in GRCh38 was absent from the variant-containing haplotype because one or both nucleotides were altered, removed or separated by the variant. Because methylation is undefined on haplotypes lacking the CpG, methylation was quantified only on haplotypes carrying the CpG-containing allele. Var-CpGs were classified as SNV-, indel- or SV-associated according to the corresponding variant class.

Var-CpG events were defined at the individual CpG level. Each SNV- or indel-associated CpG gain or loss was counted as one event, whereas each CpG affected by an SV was counted separately, allowing a single SV to contribute multiple var-CpG events. When a var-CpG intersected multiple variant records, variant-specific records were retained separately in haplotype-level analyses. For population-level analyses, duplicated records corresponding to the same var-CpG were merged into a single event.

### Graph-based interval-overlap analysis

Intersections were calculated directly between intervals represented in graph coordinates. Variant and CpG intervals were encoded as oriented graph paths with start and end offsets on their terminal segments. To reduce the search space, variants were indexed by the graph segments traversed by their paths, and each CpG interval was compared only with variants sharing at least one segment. For each candidate pair, the intervals were resolved to segment-level base positions, accounting for segment orientation. Overlap was calculated as the number of segment-base positions shared by the CpG and variant intervals. Pairs with non-zero overlap were reported together with the CpG identifier, variant identifier, overlap length and source interval annotations. This procedure enabled interval intersections to be evaluated directly in graph space without projection to a single linear reference coordinate system.

### Diversity across methylomes

For each haplotype-resolved methylome, CpG abundance was summarized as the total number of CpGs detected, and genome-wide methylation as the median methylation level across all CpGs. Summary statistics were calculated by population, continental group and sex, where applicable. Methylation levels were represented as proportions ranging from 0 to 1. Statistical analyses were performed in R v4.4.1.

To quantify between-population variability within each continental group, we first calculated the population-level median CpG abundance or methylation level and then calculated the median absolute deviation (MAD) of these population-level values within each continental group. MAD was defined as the median absolute deviation from the corresponding continental group median.

Differences in CpG abundance, CpG density and methylation levels between two groups were assessed using two-sided Wilcoxon rank-sum tests, whereas comparisons across more than two populations or continental groups used Kruskal–Wallis tests. Differences in categorical distributions, including variant and CpG classes, were assessed using Pearson’s chi-squared tests. Enrichment of genomic annotations was evaluated using Fisher’s exact tests. Unless otherwise stated, *P* values were adjusted for multiple testing using the Benjamini–Hochberg procedure^91^, with FDR < 0.05 considered significant.

### Integrated representation of CpG copy number and methylation at var-CpGs

CpG copy number was represented as a discrete variable with three states (0, 1 or 2), whereas CpG methylation was represented as a continuous value ranging from 0 to 100%. To integrate these measurements, each var-CpG observation was encoded as a vector with three elements corresponding to the three CpG copy number states.

The element corresponding to the observed copy number state was assigned the methylation value, whereas the two inactive elements were set to −1 to distinguish them from a methylation value of 0.

A convolutional encoder–decoder network transformed the three-element input into a 10-channel hidden representation, which was compressed into a one-dimensional latent value^56^. The decoder expanded this value through a 10-channel hidden layer to reconstruct the original input. Model performance was comparable between the training and test sets and across continental groups (Supplementary Fig. 1). The one-dimensional latent value accurately recovered both CpG copy number state (area under the receiver operating characteristic curve (AUROC) = 1.00) and methylation level (Pearson’s r = 0.99, *P* < 2.2 × 10⁻¹⁶). This latent value was therefore used as the integrated representation of CpG copy number and methylation in downstream analyses.

### Variance partitioning and population structure analysis

To quantify population-associated epigenomic variation, we analyzed three complementary CpG-level representations: methylation at shared CpGs, CpG copy number at var-CpGs, and an integrated var-CpG signal combining CpG copy number and methylation. Shared CpGs were defined as autosomal reference CpGs present in more than 99% of analyzed haplotypes. Var-CpGs were restricted to autosomal sites with intermediate presence frequencies (5% < AF < 95%), thereby excluding rare and near-fixed CpG gain or loss events.

For shared CpGs, the response variable was methylation level. For the CpG gain/loss model, the response variable was CpG copy number, encoded as 0, 1 or 2 copies per individual. For the integrated model, the response variable was the one-dimensional latent representation of CpG copy number and methylation generated by the encoder–decoder network described above. Analyses were performed separately using continental group and population labels as categorical predictors.

Following previous analyses of gene expression and splicing^57^, a linear model was fitted separately for each CpG or var-CpG, with the CpG-level measurement as the response and either continental group or population label as the explanatory variable. The proportion of variance associated with continental group or population labels was estimated from the model *R*^2^. Mean *R*^2^ values across CpGs or var-CpGs were used to summarize the contribution of population structure within each representation. Because the three models analyzed different biological measurements, their *R*^2^ values were interpreted within, rather than directly compared across, representations.

Statistical significance was assessed by randomly permuting continental group or population labels across individuals while preserving the CpG-level response matrices. For each of 1,000 permutations, the same CpG-level models were fitted and the mean *R*^2^ was recalculated. Empirical one-sided permutation *P* values were calculated as the proportion of permuted mean *R*^2^ values greater than or equal to the observed value.

Principal component analysis (PCA) was used to visualize population structure captured by shared-CpG methylation and var-CpG copy number. CpGs or var-CpGs with more than 10% missing values were excluded, and remaining missing values were imputed using the mean value of the corresponding CpG or var-CpG across individuals. PCA was performed on centered and scaled methylation or copy number matrices.

### Population-specific CpG and methylation analyses

Population-specific CpGs and var-CpGs were identified through pairwise comparisons between continental groups. Analyses were restricted to autosomal sites to avoid confounding by sex chromosome dosage and pseudoautosomal regions. Shared CpGs were evaluated using methylation differences, whereas var-CpGs were evaluated using two representations: CpG gain/loss and the integrated CpG copy number and methylation score.

Pairwise methylation differences at shared CpGs were tested using linear models implemented in limma^92^. CpGs with more than 10% missing values were excluded, and β-values were transformed to M-values before modeling^93^. For each comparison, M-values were modeled as a function of continental group, with sex included as a covariate, and empirical Bayes moderation was used to obtain CpG-level test statistics. Statistical testing was performed on the M-value scale, whereas effect sizes were reported as differences in median β values between groups.

Analogous limma models were fitted for var-CpGs using either CpG copy number or the integrated score scaled from 0 to 1 as the response variable. Var-CpGs with more than 10% missing values were excluded. For each pairwise comparison, the var-CpG measure was modeled as a function of continental group, with sex included as a covariate, and empirical Bayes moderation was used to obtain var-CpG-level test statistics.

Unless otherwise specified, CpGs and var-CpGs were classified as population-specific when they showed an absolute between-group difference greater than 0.20 and an FDR below 0.05. Here, the difference, Δ, represented the difference in median β value for shared-CpG methylation, the difference in the scaled CpG gain/loss measure for var-CpG copy number, or the difference in the scaled integrated score. As a sensitivity analysis, we applied the same effect size threshold together with a nominal *P* < 5.0 × 10⁻⁸.

### Enrichment of CpG-altering variants at loci under directional selection

Genomic loci showing evidence of directional selection were obtained from Akbari et al.^60^ and harmonized to the GRCh38 reference assembly using UCSC liftOver with the hg19ToHg38.over.chain.gz chain file. Loci that could not be uniquely mapped were excluded. CpG-altering variants were defined as variants that created or disrupted a CpG dinucleotide. Variants with a minor allele frequency (MAF) greater than 1% were retained for analysis.

Enrichment was assessed by comparing the proportion of CpG-altering variants among directional selection loci with that of eligible non-CpG-altering variants from the same HPRC2 variant set. Variants were matched by chromosome, position, reference allele and alternative allele after harmonization of genome builds. Enrichment was evaluated using Pearson’s chi-squared test, with a two-sided *P* value below 0.05 considered significant.

### Genetic and epigenetic differentiation analyses

For each var-CpG, genetic differentiation and differentiation of the integrated var-CpG signal were quantified by comparing each focal continental group with all remaining groups. Genetic differentiation was estimated using *F*_ST_ for the variant causing CpG gain or loss. *F*_ST_ was calculated using VCFtools v0.1.16 with the Weir and Cockerham estimator^94^. Differentiation of the integrated var-CpG signal was estimated using *P*ST, the phenotypic analogue of *F*_ST_ ^14,62^. For each var-CpG, *P*_ST_ was calculated from the integrated score combining CpG presence or absence with methylation on CpG-containing haplotypes:

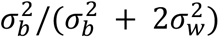

where *σ_b_*^2^ is the variance between the focal and comparison groups and *σ_w_*^2^ is the mean within-group variance. *P*_ST_ ranges from 0 to 1, with higher values indicating that a greater proportion of variation in the integrated var-CpG signal occurs between groups rather than within groups.

To identify var-CpGs for which integrated differentiation deviated from that predicted by genetic differentiation, *P*_ST_ was regressed on *F*_ST_ separately for each focal-versus-rest comparison. The fitted value represented the expected *P*_ST_ given the observed *F*_ST_. Var-CpGs were classified as having lower-than-expected integrated differentiation when expected *P*_ST_ minus observed *P*_ST_ exceeded 0.05, and higher-than-expected integrated differentiation when observed *P*_ST_ minus expected *P*_ST_ exceeded 0.05. These categories were defined using a descriptive residual threshold rather than a formal significance test.

### Association between var-CpG methylation and genetic features

Associations between var-CpG methylation M-values and genetic features were tested using linear models implemented in the MatrixEQTL R package^95^. In total, 2,816,359 intermediate-frequency var-CpGs (5% < AF < 95%) were tested. Three classes of genetic features were analyzed separately: copy number of the target var-CpG itself, copy number of nearby var-CpGs and nearby genetic variants that did not directly cause CpG gain or loss. In each model, var-CpG methylation was the response variable and the corresponding genetic feature was the predictor, with sex and the first five population principal components included as covariates. Nearby features were tested within 10 kb of the target var-CpG^17,18,96^ using coordinates projected onto the GRCh38 backbone.

Significance was assessed using FastQTL-style permutation calibration^97^. For each target var-CpG, null association statistics were generated from 1,000 permutations and used to fit a beta distribution, from which the observed nominal *P* value was converted to a beta-adjusted empirical *P* value. Multiple testing was controlled at an estimated FDR of 5% using the qvalue R package. Sensitivity analyses repeated the association tests using MAF thresholds of 5%, 10% and 20%.

### ChromHMM and regulatory-element enrichment analysis

ChromHMM annotations for GM12878 lymphoblastoid cell lines (LCLs), generated from histone modification profiles, were obtained from the ENCODE Broad ChromHMM resource^98^ (http://hgdownload.cse.ucsc.edu/goldenPath/hg19/encodeDCC/wgEncodeBroadHmm). Coordinates were converted from hg19 to GRCh38 using UCSC liftOver and the hg19ToHg38.over.chain.gz chain file. LCL super-enhancer annotations were obtained from SEdb^99^ (http://www.licpathway.net:8081/sedb/), and gene annotations were obtained from the UCSC Genome Browser^100^ (https://genome.ucsc.edu/).

To assess enrichment of regulatory annotations among var-CpGs with multilayer genetic associations, the foreground comprised var-CpGs whose methylation was significantly associated with all three genetic feature classes: self-copy number, nearby var-CpG copy number and nearby non-CpG-altering variants. The background comprised all var-CpGs whose methylation was significantly associated with at least one of these feature classes. For each ChromHMM state or regulatory element, enrichment was evaluated using Fisher’s exact test. *P* values were adjusted using the Benjamini–Hochberg procedure^91^, with FDR < 0.05 considered significant. Fold enrichment was calculated as the proportion of foreground var-CpGs overlapping an annotation divided by the corresponding proportion among background var-CpGs.

### Association between promoter var-CpG methylation and transcript expression

Associations between promoter var-CpG methylation and transcript expression were tested in 195 individuals with matched methylation, genetic-variant and Iso-Seq transcript-expression data. Transcript-specific promoters were defined as the strand-aware 2-kb interval upstream of each annotated TSS. Iso-Seq transcript-expression data were generated, integrated, filtered and inverse-normal transformed as described in the accompanying study^30^.

In total, 97,712 promoter var-CpGs associated with 165,223 annotated transcripts were evaluated. For each promoter var-CpG–transcript pair, transcript expression was modeled as a function of the var-CpG methylation M-value, with sex, the first five principal components derived from genome-wide genotypes and one expression PEER factor associated with technical batch effects included as covariates. To evaluate the contribution of the underlying CpG-altering variant, a second model additionally included CpG copy number, encoded as 0, 1 or 2 copies per individual. Associations significant both before and after adjustment for CpG copy number were classified as robust to adjustment for the underlying variant, whereas associations detected only after adjustment were classified as conditional methylation–expression associations.

Multiple testing was addressed using a two-stage hierarchical procedure^101^. First, promoter var-CpG association *P* values were combined for each transcript using the aggregated Cauchy association test^102^. Transcript-level *P* values were then adjusted across transcripts using the qvalue procedure. For transcripts with q < 0.05, promoter var-CpG-level *P* values were adjusted within each transcript using the Benjamini–Hochberg procedure^91^. Var-CpG–transcript pairs with within-transcript FDR < 0.05 were considered significant.

### Mediation analysis of genetic effects through var-CpG methylation

To test whether var-CpG methylation statistically mediated associations between CpG gain/loss and transcript expression, we fitted a mediator model in which var-CpG methylation was modeled as a function of CpG copy number and covariates, and an outcome model in which transcript expression was modeled as a function of CpG copy number, var-CpG methylation and the same covariates. CpG copy number was encoded as 0, 1 or 2 copies per individual. Both models adjusted for sex, the first five principal components derived from genome-wide genotypes and one expression PEER factor associated with technical batch effects.

Mediation was evaluated using the mediation R package^103^ with nonparametric bootstrap simulations. Analyses were initially performed with 100 simulations, and associations yielding a bootstrap *P* value of zero were rerun with 10,000 simulations. To reduce the computational burden, tests were restricted to var-CpG–transcript pairs for which CpG copy number was associated with var-CpG methylation at nominal *P* < 0.1 in the mediator model. Average causal mediation effect *P* values were adjusted using the Benjamini–Hochberg procedure^91^, and statistical mediation was considered significant at FDR < 0.05.

### Variant-by-methylation interaction analysis

Variant-by-methylation interactions were tested using linear models in which transcript expression was modeled as a function of CpG copy number, var-CpG methylation and their interaction. CpG copy number was encoded as 0, 1 or 2 copies per individual, and methylation was represented by the corresponding M-value. Models were adjusted for sex, the first five principal components derived from genome-wide genotypes and one expression PEER factor associated with technical batch effects.

Interaction analyses were restricted to var-CpG–transcript pairs showing both significant CpG copy number–expression and methylation–expression associations in the copy number-adjusted expression model (FDR < 0.05 for both). Interaction-term *P* values were adjusted using the Benjamini–Hochberg procedure^91^, and variant-by-methylation interactions were considered significant at FDR < 0.05.

### Enrichment of CpG-altering variants among molecular QTLs

Molecular QTL annotations, including mQTLs, expression QTLs (eQTLs) and splicing QTLs (sQTLs), were obtained for HPRC2 LCLs and 50 GTEx tissues^67,78,79^. HPRC2 eQTL annotations were obtained from the accompanying study^30^, whereas mQTL and sQTL annotations were obtained from the rust-fastQTL resource^78^. Enrichment analyses were restricted to CpG-altering and background variants with MAF greater than 1%. For HPRC2 LCL analyses, SNVs, indels and structural variants were included. For GTEx eQTL analyses, both CpG-altering and background variants were restricted to SNVs and indels.

For each molecular QTL class, the proportion of CpG-altering variants overlapping QTL variants was compared with the corresponding proportion among other common variants. Enrichment was evaluated using Pearson’s chi-squared test. Fold enrichment was calculated as the proportion of CpG-altering variants overlapping a given QTL class divided by the corresponding proportion among background variants.

### Clinical and pharmacogenomic annotation of CpG-altering variants

Clinical annotations were obtained from ClinVar^82^ (release 20251006; https://www.ncbi.nlm.nih.gov/clinvar/), and pharmacogenomic annotations were obtained from ClinPGx^83^ (https://www.clinpgx.org/). Variants were matched to ClinVar and ClinPGx records by chromosome, position, reference allele and alternative allele after harmonization of genome build and allele orientation.

Enrichment analyses were restricted to CpG-altering and background variants with MAF greater than 1%, unless otherwise specified. Background variants comprised other common variants from the same HPRC2 variant set^30^. For each annotation category, the proportion of CpG-altering variants carrying a clinical or pharmacogenomic annotation was compared with the corresponding proportion among background variants. ClinVar annotations were evaluated overall and by clinical-significance category, including pathogenic or likely pathogenic, benign or likely benign, risk and protective variants. Enrichment was assessed using Fisher’s exact test, and *P* values were adjusted using the Benjamini–Hochberg procedure where applicable^91^. Fold enrichment was calculated as the proportion of CpG-altering variants in a given annotation category divided by the corresponding proportion among background variants.

Promoters were defined as strand-aware 2-kb intervals upstream of TSSs using GRCh38 gene annotations obtained from the UCSC Genome Browser^100^. GRCh38 promoter and CGI annotations were used to assess whether methylation differences among clinically and pharmacogenomically annotated var-CpGs varied by genomic context. Because methylation levels showed negligible correlations with CpG frequency for both reference CpGs (Spearman’s ρ = −0.008) and non-reference CpGs (Spearman’s ρ = −0.011), these comparisons were not further stratified by CpG frequency (Supplementary Fig. 2).

Population-specific clinical and pharmacogenomic var-CpGs were identified by intersecting annotated CpG-altering variants with population-specific var-CpGs detected in pairwise comparisons between continental groups. These analyses were restricted to variants with MAF greater than 5% and were performed using both the CpG gain/loss and integrated models.

## Data availability

Metadata and download URIs available through the HPRC Data Explorer on https://humanpangenome.org/. Assemblies and select annotations are available through a UCSC Genome Browser Hub (https://hgdownload.gi.ucsc.edu/hubs/HPRC/index.html) as well as the Ensembl Genome Browser (https://projects.ensembl.org/hprc/)^104^. Assembly and epigenomic data is available in the HPRC Epigenome Browser (https://epigenome.humanpangenome.org/). Haplotype-based genetic variants were downloaded from https://s3-us-west-2.amazonaws.com/human-pangenomics/index.html?prefix=submissions/759B21AD-0ED8-4640-A433-7C92A57EA3D3--UW_EEE_SV_Calls/GRCh38/. Graph-based genetic variants were downloaded from https://github.com/human-pangenomics/hpp_pangenome_resources. Haplotype-resolved genome-assembly annotations are available at https://github.com/human-pangenomics/hprc_intermediate_assembly/tree/main/data_tables/annotation.

## Code availability

Code used for the analyses presented in this study is available at https://github.com/twlab/Panepigenome.

## Acknowledgements

We thank colleagues in the Human Pagenome Reference Consortium for helpful discussions and feedback. This work was supported by NIH grants U41HG010972, U41HG010971, U24HG012070, U24NS132103, and UM1DA058219.

## Author Contributions

T.W. and Z.D. conceived the study. Z.D. designed the analytical framework and performed the principal computational and statistical analyses, including panepigenome construction, graph coordinate projection, CpG and var-CpG annotation, population epigenomic analyses, structural variant and transposable element analyses, transcript association and mediation analyses, molecular QTL enrichment analyses, and clinical and pharmacogenomic analyses. Z.D. interpreted the results, prepared the figures and wrote the original draft. R.S.F. contributed to DNA methylation data generation. Z.D., C.T. and J.F.M. contributed to DNA methylation quality control. W.Z. developed tools for integrating DNA methylation data into graph coordinates. S.R.J. contributed to the development of the integrated model using a convolutional neural network with an encoder–decoder architecture. E.A.B., R.L., J.F.M., X.Z. and Z.D. contributed to phasing, alignment and methylation profiling. T.W. and Z.D. revised the manuscript. J.F.M., X.Z., J.J., S.D., T.L. and D.L. reviewed and edited the manuscript. T.W. supervised the project and acquired funding. All authors reviewed and approved the final manuscript.

## Declaration of generative AI and AI-assisted technologies in scientific writing

During the preparation of the manuscript, the authors used ChatGPT to improve language and readability. After using this tool, the authors reviewed and edited the content as needed and take full responsibility for the content of the publication.

## Human Pangenome Reference Consortium Version 2

### Authors

Derek Albracht^1^, Ivan A. Alexandrov^2^, Jamie Allen^3^, Alawi A. Alsheikh-Ali^4^, Nicolas Altemose^5^, Casey Andrews^6^, Dmitry Antipov^7^, Lucinda Antonacci-Fulton^1^, Alexander Arguello^8^, Mobin Asri^9^, Marcelo Ayllon^10^, Jennifer R. Balacco^11^, Floris P. Barthel^12^, Edward A. Belter Jr^1^, Halle D. Bender^9^, Andrew P. Blair^9^, Davide Bolognini^13^, Katherine E. Bonini^14^, Christina Boucher^15^, Guillaume Bourque^16,17,18^, Silvia Buonaiuto^19^, Shuo Cao^19^, Andrew Carroll^20^, Ann M. Mc Cartney^21^, Monika Cechova^9^, Mark J.P. Chaisson^22^, Pi-Chuan Chang^20^, Xian Chang^9^, Jitender Cheema^3^, Haoyu Cheng^23^, Claudio Ciofi^24^, Hiram Clawson^9^, Sarah Cody^1^, Vincenza Colonna^19^, Holland C. Conwell^25^, Robert Cook-Deegan^26^, Mark Diekhans^9^, Maria Angela Diroma^24^, Daniel Doerr^27,28,29^, Zheng Dong^6^, Danilo Dubocanin^5^, Richard Durbin^30,31^, Jana Ebler^27,32^, Evan E. Eichler^10,33^, Jordan M. Eizenga^9^, Parsa Eskandar^9^, Eddie Ferro^15^, Anna-Sophie Fiston-Lavier^34,35^, Sarah M. Ford^25^, Willard W. Ford^36^, Giulio Formenti^11^, Adam Frankish^3^, Mallory A. Freeberg^3^, Qichen Fu^6^, Stephanie M. Fullerton^37^, Robert S. Fulton^1^, Shenghan Gao^38^, Yan Gao^39^, Gage H. Garcia^10^, Obed A. Garcia^40^, Joshua M.V. Gardner^9^, Shilpa Garg^41^, Erik Garrison^19^, Nanibaa’ A. Garrison^42,43,44^, John E. Garza^1^, Margarita Geleta^45,46^, Mohammadmersad Ghorbani^47^, Tina A. Graves-Lindsay^1^, Richard E. Green^25^, Carol W. Greider^48^, Cristian Groza^49^, Bida Gu^22^, Andrea Guarracino^12,19^, Melissa Gymrek^50^, Maximilian Haeussler^9^, Leanne Haggerty^3^, Ira M. Hall^51,52^, Nancy F. Hansen^7^, Yue Hao^12^, Mohammad Amiruddin Hashmi^4^, David Haussler^9^, Prajna Hebbar^9^, Peter Heringer^27,28,29^, Glenn Hickey^9^, Todd L. Hillaker^9^, S. Nakib Hossain^3^, Neng Huang^39,53^, Sarah E. Hunt^3^, Toby Hunt^3^, Alexander G. Ioannidis^5,9,46^, Nafiseh Jafarzadeh^9^, Nivesh Jain^11^, Erich D. Jarvis^11,33^, Maryam Jehangir^12^, Juan Jiang^6^, Eimear E. Kenny^14^, Juhyun Kim^7^, Bonhwang Koo^11^, Sergey Koren^7^, Milinn Kremitzki^1,6^, Charles H. Langley^54^, Ben Langmead^55^, Heather A. Lawson^6^, Daofeng Li^6^, Heng Li^39,53^, Ronghan Li^6^, Wen-Wei Liao^51,52^, Jiadong Lin^10^, Tianjie Liu^6^, Glennis A. Logsdon^38^, Ryan Lorig-Roach^9^, Jonathan LoTempio Jr^21,56^, Hailey Loucks^9^, Jane E. Loveland^3^, Jianguo Lu^57^, Shuangjia Lu^51,52^, Julian K. Lucas^9^, Walfred Ma^22^, Juan F. Macias-Velasco^1,6,58^, Kateryna D. Makova^59^, Maximillian G. Marin^39,53^, Christopher Markovic^1^, Tobias Marschall^27,32^, Franco L. Marsico^19^, Fergal J. Martin^3^, Mira Mastoras^9^, Capucine Mayoud^34^, Brandy McNulty^9^, Jack A. Medico^11^, Julian M. Menendez^9^, Karen H. Miga^9^, Anna Minkina^60^, Matthew W. Mitchell^61^, Saswat K. Mohanty^62^, Younes Mokrab^47,63,64^, Jean Monlong^65^, Shabir Moosa^47^, Avelina Moreno-Ochando^66,67^, Shinichi Morishita^68^, Jonathan M. Mudge^3^, Katherine M. Munson^10^, Njagi Mwaniki^69^, Nasna Nassir^4^, Chiara Natali^24^, Shloka Negi^9^, Lingbin Ni^10^, Adam M. Novak^9^, Faith Okamoto^9^, Keisuke K. Oshima^38^, Pilar N. Ossorio^70,71^, Chie Owa^68^, Sadye Paez^11^, Benedict Paten^9^, Clelia Peano^13,72^, Adam M. Phillippy^7,55,73,74^, Brandon D. Pickett^7^, Laura Pignata^19^, Nadia Pisanti^69^, David Porubsky^10,75^, Pjotr Prins^19^, Timofey Prodanov^27,32^, Anandi Radhakrishnan^9^, T. Rhyker Ranallo-Benavidez^12^, Brian J. Raney^9^, Mikko Rautiainen^76^, Alessandro Raveane^13^, Andreas Rechtsteiner^48^, Luyao Ren^10,33^, Arang Rhie^7^, Fedor Ryabov^77,78^, Samuel Sacco^25^, Farnaz Salehi^19^, Michael C. Schatz^55,79^, Laura B. Scheinfeldt^80^, Aarushi Sehgal^36^, William E. Seligmann^25^, Mahsa Shabani^81^, Kishwar Shafin^20^, Shadi Shahatit^34^, Ruhollah Shemirani^14^, Vikram S. Shivakumar^55^, Swati Sinha^3^, Jouni Sirén^9^, Linnéa Smeds^62^, Steven J. Solar^7^, Marco Sollitto^11,24^, Nicole Soranzo^13,30,82^, Andrew B. Stergachis^10,60^, Marie-Marthe Suner^3^, Yoshihiko Suzuki^68^, Arda Söylev^27,32^, Ahmad Abou Tayoun^83,84^, Jack A.S. Tierney^3^, Chad Tomlinson^1^, Francesca Floriana Tricomi^3^, Mohammed Uddin^4,85^, Matteo Tommaso Ungaro^25,86^, Rahul Varki^15^, Flavia Villani^19^, Ivo Violich^9^, Mitchell R. Vollger^87^, Brian P. Walenz^7^, Charles Wang^88^, Lisa E. Wang^14^, Ting Wang^1,6,58^, Aaron M. Wenger^89^, Conor V. Whelan^11^, Zilan Xin^6^, Zheng Xu^6^, Kai Ye^90^, DongAhn Yoo^10^, Wenjin Zhang^6^, Ying Zhou^39^, Xiaoyu Zhuo^6^, Giulia Zunino^13^

### Affiliations

^1^ McDonnell Genome Institute, Washington University School of Medicine, St. Louis, MO 63108, USA

^2^ Department of Human Molecular Genetics and Biochemistry, Faculty of Medical and Health Sciences, Tel Aviv University, Tel Aviv 69978, Israel

^3^ European Molecular Biology Laboratory, European Bioinformatics Institute (EMBL-EBI), Wellcome Genome Campus, Hinxton, Cambridge CB10 1SD, UK

^4^ Center for Applied and Translational Genomics (CATG), Mohammed Bin Rashid University of Medicine and Health Sciences, Dubai Health, Dubai, UAE

^5^ Department of Genetics, Stanford University, Palo Alto, CA 94304 USA

^6^ Department of Genetics, Washington University School of Medicine, St. Louis, MO 63110, USA

^7^ Genome Informatics Section, Center for Genomics and Data Science Research, National Human Genome Research Institute, National Institutes of Health, Bethesda, MD 20892, USA

^8^ Division of Genome Sciences, National Human Genome Research Institute, Bethesda, MD 20871 USA

^9^ UC Santa Cruz Genomics Institute, University of California, Santa Cruz, CA 95060, USA

^10^ Department of Genome Sciences, University of Washington School of Medicine, Seattle, WA 98195, USA

^11^ The Vertebrate Genome Laboratory, The Rockefeller University, New York, NY 10065, USA

^12^ Bioinnovation and Genome Sciences, The Translational Genomics Research Institute (TGen), Phoenix, AZ 85004, USA

^13^ Human Technopole, Milan, Italy

^14^ Institute for Genomic Health, Icahn School of Medicine at Mount Sinai, New York, NY 10029, USA

^15^ Department of Computer and Information Science and Engineering, University of Florida, Gainesville, FL 32611, USA

^16^ Canadian Center for Computational Genomics, McGill University, Montréal, QC H3A 0G1, Canada

^17^ Department of Human Genetics, McGill University, Montréal, QC H3A 0G1, Canada

^18^ Victor Phillip Dahdaleh Institute of Genomic Medicine, Montréal, QC H3A 0G1, Canada

^19^ Department of Genetics, Genomics and Informatics, University of Tennessee Health Science Center, Memphis, TN 38163, USA

^20^ Google LLC, Mountain View, CA 94043, USA

^21^ Institute of Clinical and Translational Sciences, University of California, Irvine, CA 92697, USA

^22^ Quantitative and Computational Biology, University of Southern California, Los Angeles, CA 90089, USA

^23^ Department of Biomedical Informatics and Data Science, Yale School of Medicine, New Haven, CT 06510, USA

^24^ Department of Biology, University of Florence, Sesto Fiorentino, FI 50019, Italy

^25^ Department of Ecology and Evolutionary Biology, University of California, Santa Cruz, CA 95060, USA

^26^ Arizona State University, Consortium for Science, Policy & Outcomes, Washington, DC 20006, USA

^27^ Center for Digital Medicine, Heinrich Heine University Düsseldorf, Düsseldorf, NRW, DE

^28^ Department for Endocrinology and Diabetology at the Medical Faculty and University Hospital Düsseldorf, Heinrich Heine University Düsseldorf, Düsseldorf, NRW, DE

^29^ Paul-Langerhans-Group Computational Diabetology, German Diabetes Center (DDZ) and Leibniz Institute for Diabetes Research, Düsseldorf, NRW, DE

^30^ Wellcome Sanger Institute, Genome Campus, Hinxton, CB10 1RQ, UK

^31^ Department of Genetics, University of Cambridge, Cambridge, CB2 3EH, UK

^32^ Institute for Medical Biometry and Bioinformatics, Medical Faculty and University Hospital Düsseldorf, Heinrich Heine University, Düsseldorf, NRW, DE

^33^ Howard Hughes Medical Institute, Chevy Chase, MD 20815, USA

^34^ ISEM, Univ Montpellier, CNRS, IRD, Montpellier, FR

^35^ Institut Universitaire de France, Paris, FR

^36^ Department of Computer Science and Engineering, University of California San Diego, La Jolla, CA 92093, USA

^37^ Department of Bioethics & Humanities, University of Washington School of Medicine, Seattle, WA 98195, USA

^38^ Department of Genetics, Epigenetics Institute, Perelman School of Medicine, University of Pennsylvania, Philadelphia, PA 19104, USA

^39^ Department of Data Science, Dana-Farber Cancer Institute, Boston, MA 02215, USA

^40^ Department of Anthropology, University of Kansas, Lawrence, KS 66045, USA

^41^ School of Health Sciences, University of Manchester, Manchester M13 9PL, UK

^42^ Traditional, ancestral and unceded territory of the Gabrielino/Tongva peoples, Institute for Society & Genetics, University of California, Los Angeles, Los Angeles, CA 90095, USA

^43^ Traditional, ancestral and unceded territory of the Gabrielino/Tongva peoples, Institute for Precision Health, David Geffen School of Medicine, University of California, Los Angeles, Los Angeles, CA 90095, USA

^44^ Traditional, ancestral and unceded territory of the Gabrielino/Tongva peoples, Division of General Internal Medicine & Health Services Research, David Geffen School of Medicine, University of California, Los Angeles, Los Angeles, CA 90095, USA

^45^ Department of Electrical Engineering and Computer Science, University of California, Berkeley, Berkeley, CA 94720, USA

^46^ Department of Biomedical Data Science, Stanford University School of Medicine, Stanford, CA 94305, USA

^47^ Medical and Population Genomics Lab, Sidra Medicine, Doha, Qatar

^48^ Department of Molecular Cell and Developmental Biology, University of California, Santa Cruz, CA, USA

^49^ Montreal Heart Institute, Montréal, QC, Canada

^50^ Department of Pediatrics, University of California San Diego, La Jolla, CA 92093, USA

^51^ Center for Genomic Health, Yale University School of Medicine, New Haven, CT 06510, USA

^52^ Department of Genetics, Yale University School of Medicine, New Haven, CT 06510, USA

^53^ Department of Biomedical Informatics, Harvard Medical School, Boston, MA 02115, USA

^54^ Department of Evolution and Ecology and the Center for Population Biology, University of California, One Shields, Davis, CA 95616, USA

^55^ Department of Computer Science, Johns Hopkins University, Baltimore, MD 21218, USA

^56^ Department of Pediatrics, Division of Genetics, School of Medicine, University of California, Irvine, CA 92697, USA

^57^ Sun Yat-sen University, Guangzhou, China

^58^ Edison Family Center for Genome Sciences & Systems Biology, Washington University School of Medicine, St. Louis, MO 63110, USA

^59^ Department of Biology and Center for Medical Genomics, Penn State University, University Park, PA 16802, USA

^60^ Division of Medical Genetics, Department of Medicine, University of Washington School of Medicine, Seattle, WA 98195, USA

^61^ The Jackson Laboratory for Genomic Medicine, Farmington, CT 06032, USA

^62^ Department of Biology, Penn State University, University Park, PA 16802, USA

^63^ Department of Biomedical Science, College of Health Sciences, Qatar University, Doha, Qatar

^64^ Department of Genetic Medicine, Weill Cornell Medicine-Qatar, Doha, Qatar

^65^ IRSD - Digestive Health Research Institute, University of Toulouse, INSERM, INRAE, ENVT, UPS, Toulouse, FR

^66^ MATCH biosystems, S.L., Elche, Spain

^67^ Universidad Miguel Hernández de Elche, Elche, Spain

^68^ Department of Computational Biology and Medical Sciences, The University of Tokyo, Kashiwa, Chiba 277-8561, Japan

^69^ Department of Computer Science, University of Pisa, Pisa, Italy

^70^ Law School, University of Wisconsin-Madison, Madison, WI 53706, USA

^71^ Morgridge Institute for Research, Madison, WI 53715, USA

^72^ Institute of Genetics and Biomedical Research, UoS of Milan, National Research Council, Milan, Italy

^73^ Department of Biomedical Engineering, Johns Hopkins University, Baltimore, MD 21218, USA

^74^ Department of Genetic Medicine, Johns Hopkins University School of Medicine, Baltimore, MD 21205, USA

^75^ Genome Biology Unit, European Molecular Biology Laboratory (EMBL), Heidelberg, DE

^76^ Institute for Molecular Medicine Finland, Helsinki Institute of Life Science, University of Helsinki, Helsinki, Finland

^77^ The Center for Bio- and Medical Technologies, Moscow, RUS

^78^ Centre for Biomedical Research and Technology, HSE University, Moscow, RUS

^79^ Department of Biology, Johns Hopkins University, Baltimore, MD 21218, USA

^80^ Coriell Institute for Medical Research, Camden, NJ 08103, USA

^81^ University of Amsterdam, Amsterdam, Netherlands

^82^ School of Clinical Medicine, University of Cambridge, Cambridge, CB2 0SP, UK

^83^ Center for Genomic Discovery, Mohammed Bin Rashid University, Dubai Health, UAE

^84^ Dubai Health Genomic Medicine Center, Dubai Health, UAE

^85^ GenomeArc Inc, Mississauga, ON, Canada

^86^ Department of Biology and Biotechnologies “Charles Darwin”, University of Rome “La Sapienza”, Rome 00185, IT

^87^ Department of Human Genetics and Utah Center for Genetic Discovery, University of Utah, Salt Lake City, UT, USA

^88^ Center for Genomics, Loma Linda University School of Medicine, Loma Linda, CA 92350, USA

^89^ PacBio, Menlo Park, CA 94025, USA

^90^ The first affiliated hospital of Xi’an Jiaotong University, Xi’an Jiaotong University, Xi’an, Shaanxi, 710049, China

