## Supplemental Data 1 for "Human panepigenome represents epigenomic diversity"

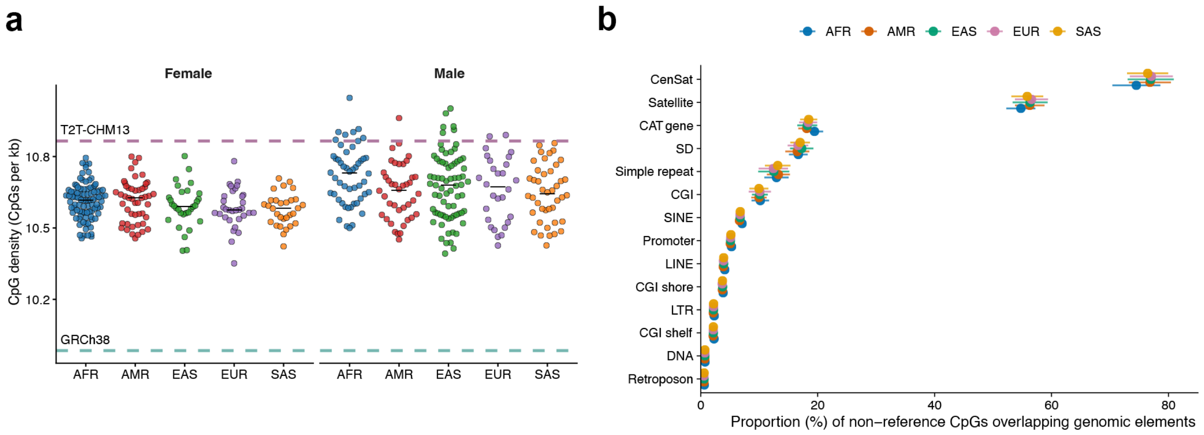


**Extended Data Fig. 1: Non-reference CpGs expand CpG-rich functional annotations.**

**a**, CpG density across 462 haplotype-resolved assemblies, GRCh38 and T2T-CHM13. CpG density was calculated as the number of CpGs per kilobase. Each point represents one haplotype-resolved assembly; teal and purple dashed horizontal lines indicate CpG densities in GRCh38 and T2T-CHM13, respectively.

**b**, Distribution of non-reference CpGs across genomic annotations and continental groups. Points indicate the mean proportion of non-reference CpGs overlapping each genomic element across assemblies within each continental group, and horizontal bars indicate one standard deviation. Genomic elements are ordered by their overall mean overlap proportion across continental groups. CenSat, centromeric satellite; CAT, Comparative Annotation Toolkit; SD, segmental duplication; CGI, CpG island; SINE, short interspersed nuclear element; LINE, long interspersed nuclear element; LTR, long terminal repeat; DNA, DNA transposon.

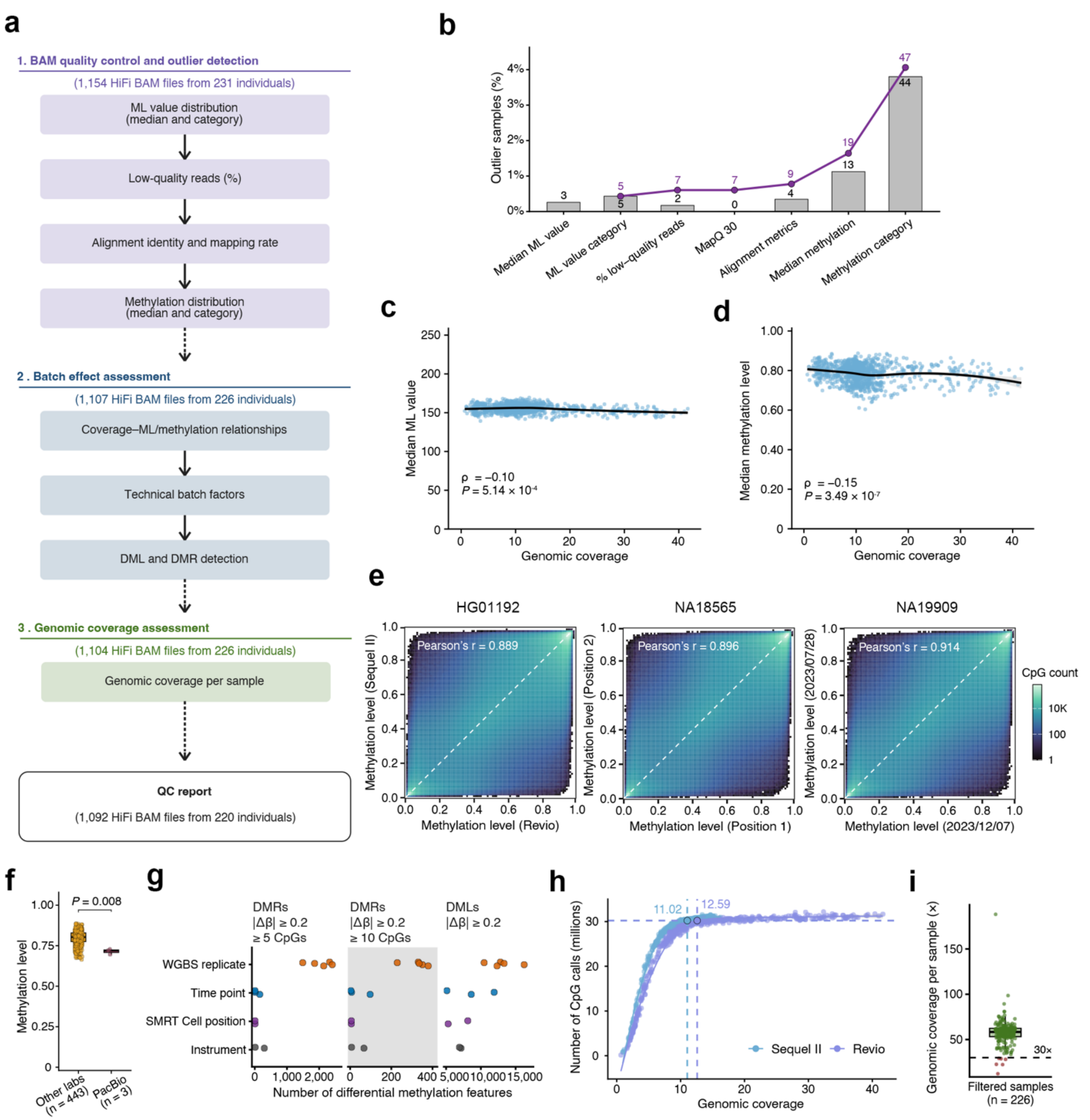


#### Extended Data Fig. 2: Sample quality control, filtering and technical validation.

**a**, Overview of the quality control framework for HiFi long-read methylation data.

**b**, Sequential identification of HiFi quality control outliers. The median ML value denotes the median base modification probability encoded by the ML tag across reads in each PacBio HiFi BAM file. Bars show the percentage of BAM files identified as outliers at each quality control step, and the purple line shows the cumulative percentage excluded across sequential filters. Numbers indicate the corresponding BAM file counts.

**c**, Relationship between genomic coverage and median ML value across BAM files. Points represent individual BAM files. The black curve shows a LOESS fit, with shading indicating the 95% confidence interval. Spearman's ρ and *P* value are shown.

**d**, Relationship between genomic coverage and median methylation level across BAM files. Points represent individual BAM files. The black curve shows a LOESS fit, with shading indicating the 95% confidence interval. Spearman's ρ and *P* value are shown.

**e**, Correlation of DNA methylation levels between technical replicate pairs across sequencing instruments (left), SMRT Cell positions (middle) and sequencing time points (right). HG01192, NA18565 and NA19909 are shown as representative examples of comparisons across sequencing instruments, SMRT Cell positions and sequencing time points, respectively. Each point represents a CpG shared between paired technical replicate BAM files from the same sample. Colors indicate point density, and dashed lines indicate the line of identity (y = x). Pearson's r is shown for each comparison.

**f**, Comparison of median methylation levels across sequencing centers. Box plots show median methylation levels for individual HiFi Revio BAM files, comparing three HG002 files generated by PacBio (m84005_220827_014912_s1.hifi_reads.bam, m84005_220919_232112_s2.hifi_reads.bam and m84011_220902_175841_s1.hifi_reads.bam) with Revio files generated by other laboratories. The PacBio HG002 files showed lower median methylation levels than files from other laboratories (72.0% versus 80.3%; Wilcoxon rank-sum test, *P* = 0.008). Points represent individual BAM files.

**g**, Technical reproducibility of DNA methylation measurements. Technical replicate comparisons were performed for seven representative samples across sequencing instruments, SMRT Cell positions and sequencing time points, with separate analyses of technical replicates from five LCL whole-genome bisulfite sequencing (WGBS) datasets. Dot plots show the numbers of differentially methylated regions (DMRs) and differentially methylated loci (DMLs) detected in each replicate comparison using the DSS R package. Points represent individual replicate pairs grouped by technical factor. DMRs were defined using $|\Delta\beta| \geq 0.20$ and either at least 5 or at least 10 CpGs; DMLs were defined using $|\Delta\beta| \geq 0.20$. WGBS replicate comparisons showed greater variability than comparisons across sequencing instruments, SMRT Cell positions or sequencing time points.

**h**, Relationship between genomic coverage and CpG call count, stratified by sequencing instrument. Points represent individual BAM files generated using the Sequel II or Revio HiFi platforms. Curves show LOESS fits. Dashed lines indicate the estimated saturation point, defined as the first coverage value at which the marginal increase in predicted CpG call count fell below 1% of the maximum observed CpG call count per unit increase in coverage.

**i**, Genomic coverage of 226 filtered samples. BAM files were merged by sample to estimate genome-wide coverage. Six samples at or below the 30× coverage threshold were excluded from downstream analyses and are shown in dark red.

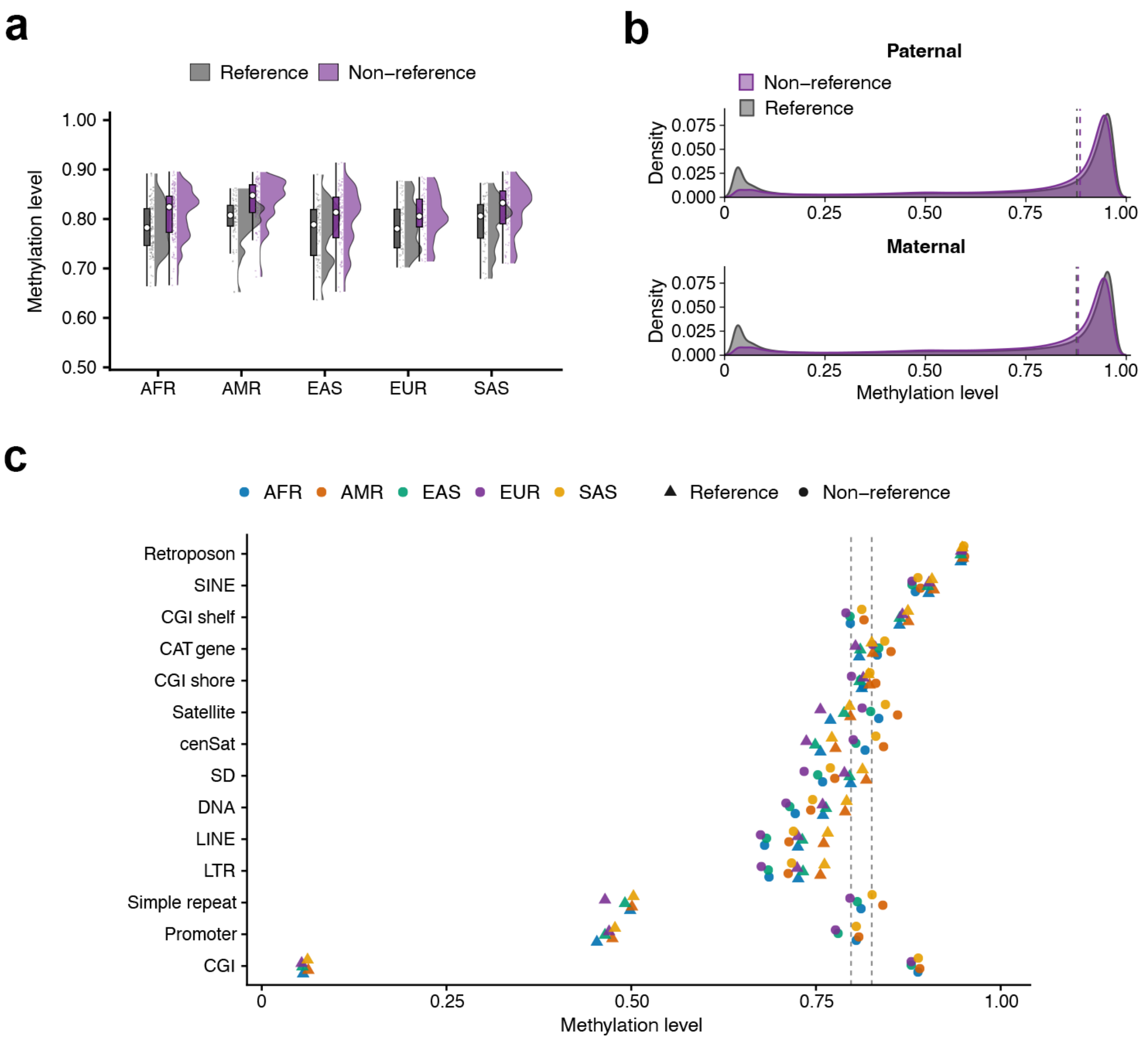


#### Extended Data Fig. 3: Non-reference CpGs show distinct methylation patterns compared with reference CpGs.

**a**, Methylation levels at non-reference and reference CpGs. Each point represents one haplotype-resolved assembly. Non-reference CpGs showed higher median methylation than reference CpGs (81.5% versus 78.8%; Wilcoxon signed-rank test, $P = 5.5 \times 10^{-16}$).

**b**, Representative haplotype-resolved methylation distributions of reference and non-reference CpGs in HG002.

**c**, Methylation levels of reference and non-reference CpGs across genomic elements. Each point represents the median DNA methylation level for reference or non-reference CpGs within a genomic element for one continental group (AFR, AMR, EAS, EUR or SAS). Colors denote continental groups, and symbol shapes distinguish reference and non-reference CpGs. Vertical dashed lines indicate the overall median methylation levels of reference CpGs (79.75%) and non-reference CpGs (82.55%) across all genomic elements. Median methylation levels were calculated separately for each genomic element and continental group. CenSat, centromeric satellite; CAT, Comparative Annotation Toolkit; SD, segmental duplication; CGI, CpG island; SINE, short interspersed nuclear element; LINE, long interspersed nuclear element; LTR, long terminal repeat; DNA, DNA transposon.

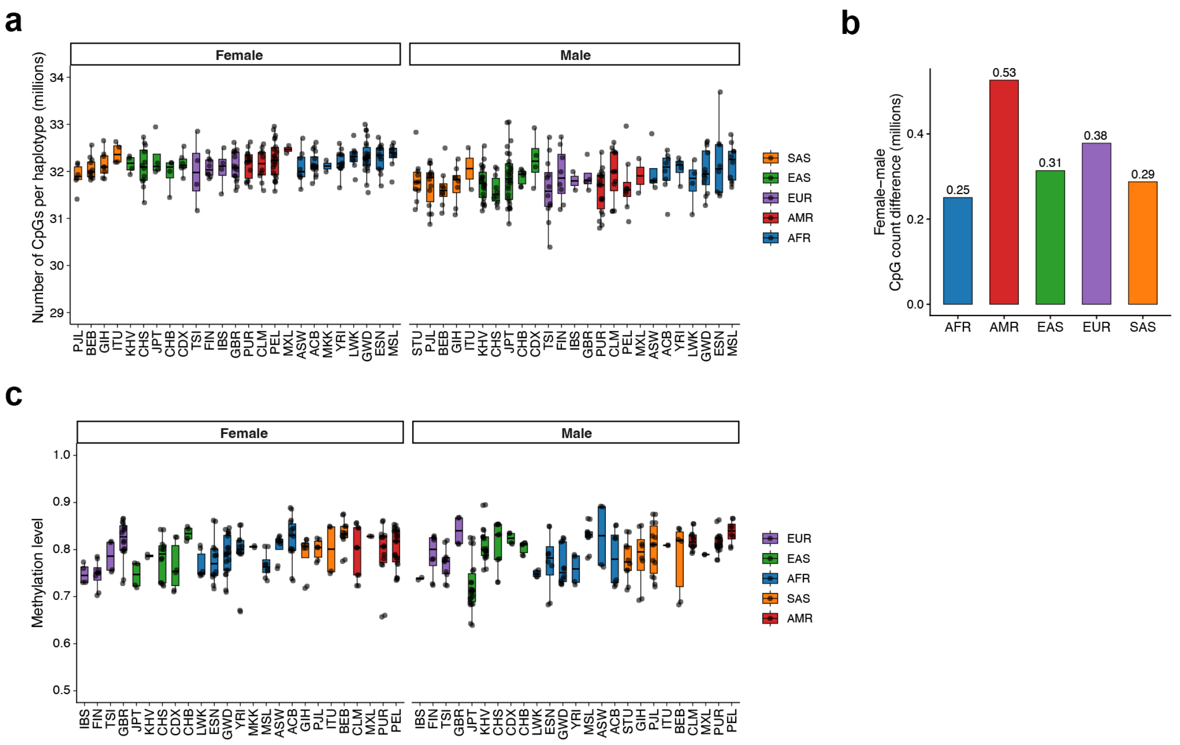


### Extended Data Fig. 4: Diversity across haplotype-resolved human methylomes.

**a**, Number of CpG sites per haplotype, stratified by population and sex. Each point represents one haplotype-resolved assembly and is colored by continental group. HG002 and HG005 were excluded because their population labels were unavailable or ambiguous. NA21309 and HG06807 were excluded because their populations were represented by only one sample each.

**b**, Sex-associated differences in assembly-derived CpG counts across continental groups.

**c**, Population- and sex-associated variation in DNA methylation. HG002 and HG005 were excluded because their population labels were unavailable or ambiguous. NA21309 and HG06807 were excluded because their populations were represented by only one sample each.

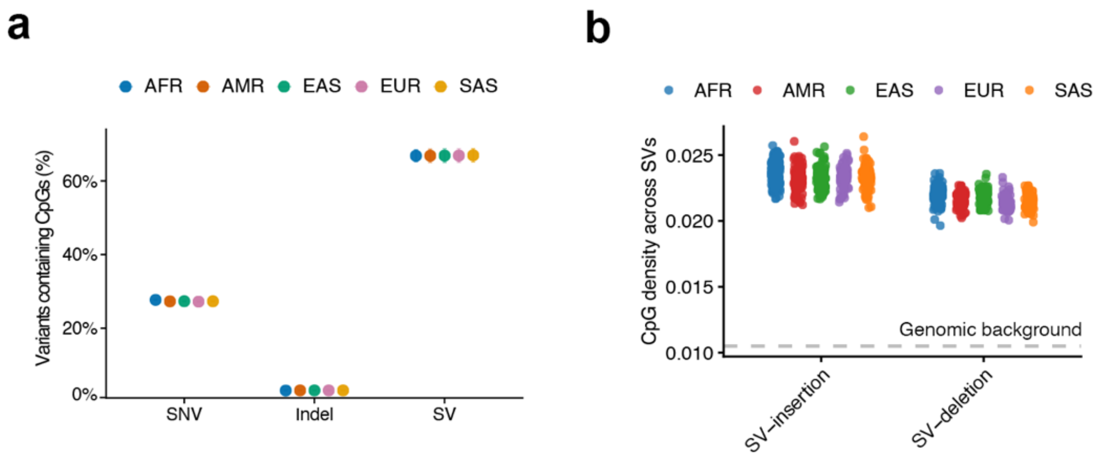


#### Extended Data Fig. 5: CpG alterations across genetic variant classes.

**a**, Mean proportion of variants containing at least one CpG across haplotype-resolved methylomes, stratified by continental group and variant class. Points indicate means, error bars indicate standard deviation, and colors denote continental groups.

**b**, CpG density within SVs. CpG density was calculated as the total number of CpGs divided by the total length of SVs. The gray dashed horizontal line indicates the mean genome-wide CpG density across HPRC2 assemblies. For each assembly, genome-wide CpG density was calculated as the total number of CpGs divided by the assembly length.

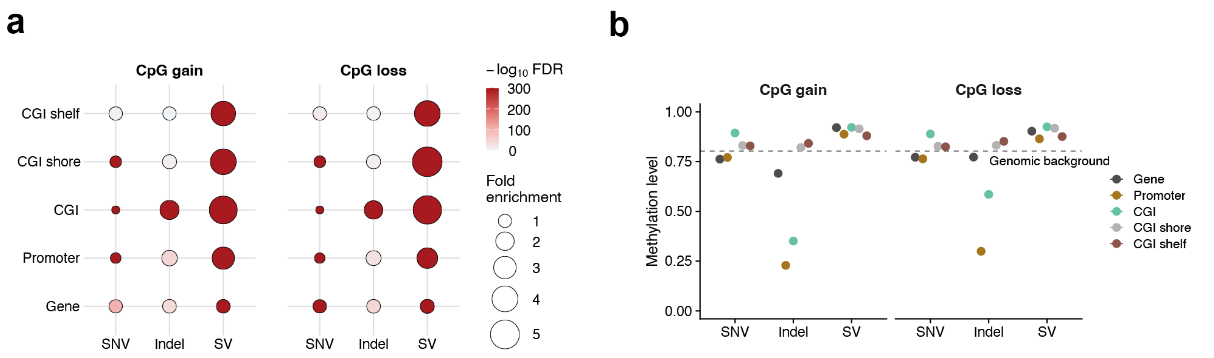


#### Extended Data Fig. 6: Genomic context of var-CpGs across haplotype-resolved methylomes.

**a**, Enrichment of CpG gain and loss events across genomic features, stratified by variant class. Bubble size indicates fold enrichment, calculated as the proportion of var-CpGs overlapping each genomic feature relative to the corresponding proportion of all CpGs overlapping that feature. Bubble color indicates $-\log_{10}$(FDR), where FDR values were obtained by Benjamini–Hochberg correction of Fisher's exact test *P* values. CGI, CpG island.

**b**, Methylation patterns of var-CpGs across genomic features. Points show median methylation levels of CpG gain and loss sites, stratified by variant class and genomic feature, and error bars indicate one standard deviation across methylomes. The dashed horizontal line indicates the genome-wide median methylation level across haplotype-resolved methylomes (80.3%). CpG gain and loss sites are shown separately for SNVs, indels and SVs. Genomic features include gene bodies, promoters, CGIs, CGI shores and CGI shelves.

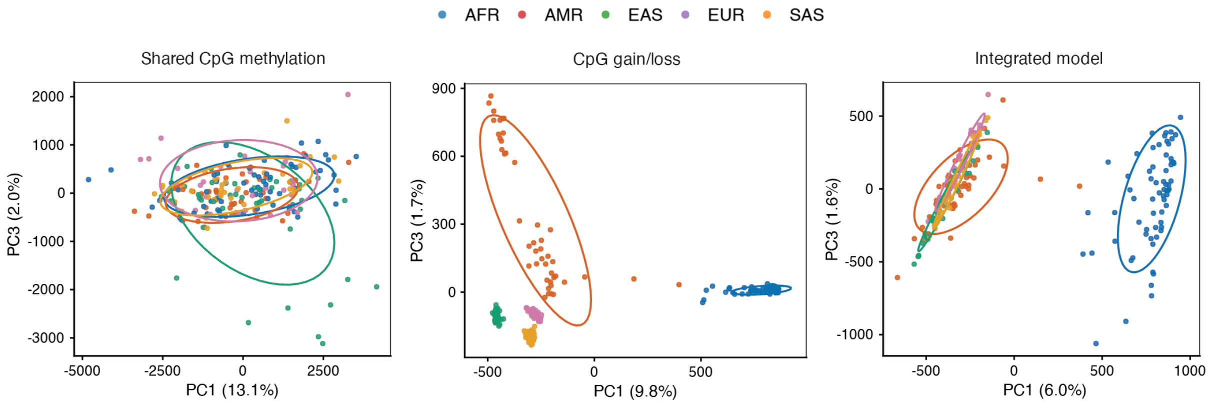


**Extended Data Fig. 7: Population structure captured by the three panepigenomic models.**

Sample loadings on PC1 and PC3 are shown for each model and are colored by continental group. From left to right, the panels represent methylation differences at shared CpGs, CpG gain and loss at var-CpGs, and an integrated model combining CpG presence or absence with methylation differences on CpG-containing haplotypes.

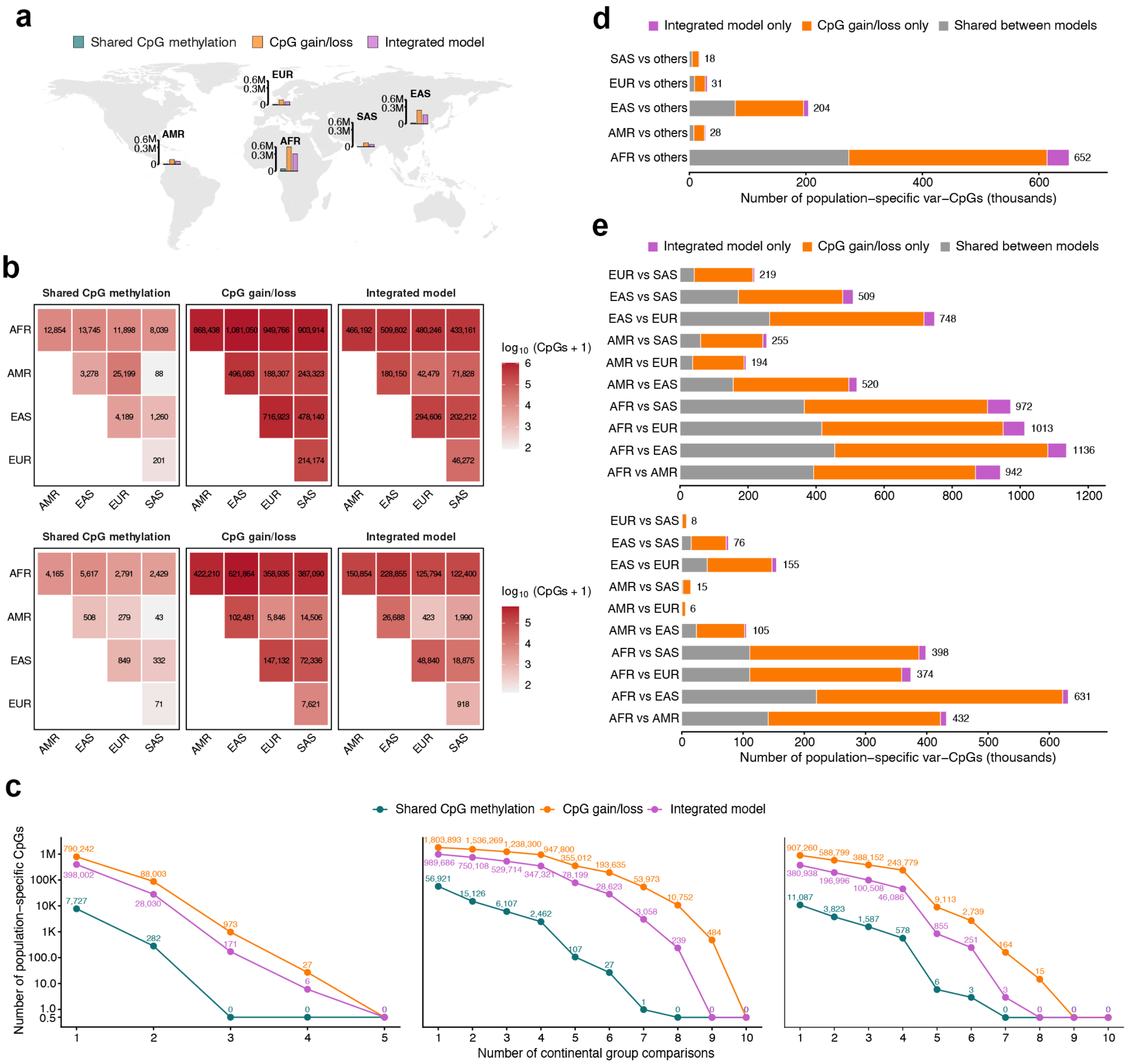


#### Extended Data Fig. 8: Var-CpGs substantially expand the catalog of population-specific CpGs.

**a**, Numbers of population-specific CpGs identified in focal-versus-rest comparisons for each continental group using $|\Delta| > 0.20$ and nominal $P < 5.0 \times 10^{-8}$.

**b**, Numbers of population-specific CpGs identified in pairwise comparisons between continental groups using either $|\Delta| > 0.20$ and FDR $< 0.05$ (upper) or $|\Delta| > 0.20$ and nominal $P < 5.0 \times 10^{-8}$ (lower).

**c**, Numbers of population-specific CpGs identified in one or more continental group comparisons for shared CpG methylation, CpG gain/loss and the integrated model. Results are shown for focal-versus-rest comparisons using $|\Delta| > 0.20$ and nominal $P < 5.0 \times 10^{-8}$ (left), pairwise comparisons using $|\Delta| > 0.20$ and FDR < 0.05 (middle), and pairwise comparisons using $|\Delta| > 0.20$ and nominal $P < 5.0 \times 10^{-8}$ (right). Points indicate the numbers of CpGs identified in the indicated numbers of continental groups. The y axis is shown on a log10 scale.

**d**, Overlap between the CpG gain/loss model and the integrated model among population-specific var-CpGs identified in focal-versus-rest comparisons, shown separately for each continental group. Population-specific var-CpGs were defined using $|\Delta| > 0.20$ and nominal $P < 5.0 \times 10^{-8}$.

**e**, Overlap between the CpG gain/loss model and the integrated model among population-specific var-CpGs identified in each pairwise comparison between continental groups. Population-specific var-CpGs were defined using $|\Delta| > 0.20$ and FDR < 0.05 (upper) or $|\Delta| > 0.20$ and nominal $P < 5.0 \times 10^{-8}$ (lower).

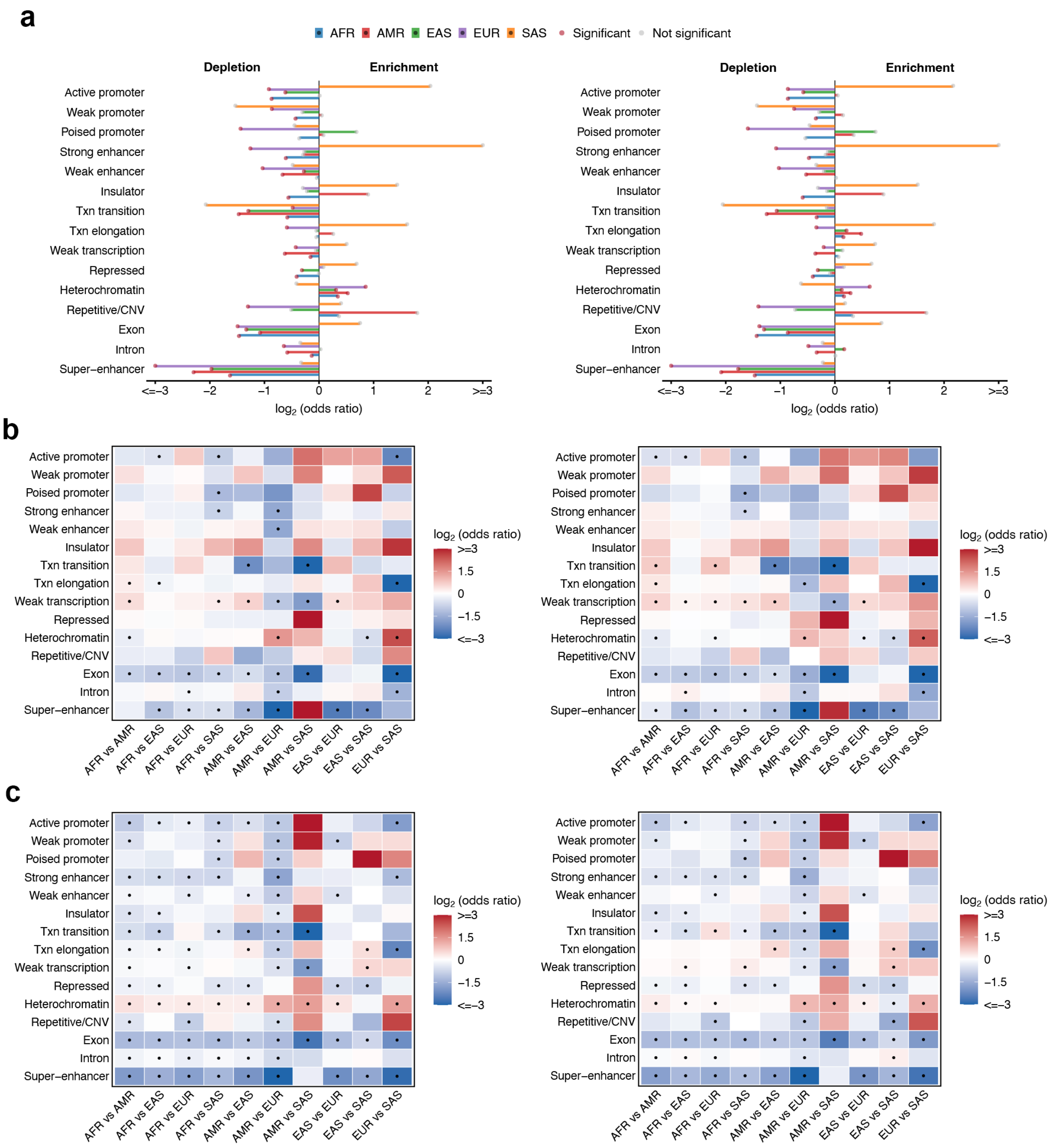


### Extended Data Fig. 9: Genomic enrichment of population-specific var-CpGs relative to population-specific shared CpGs.

**a**, Enrichment of population-specific var-CpGs across genomic annotations relative to population-specific shared CpGs in focal-versus-rest comparisons using $|\Delta| > 0.20$ and FDR $< 0.05$. Bars show the $\log_2$ odds ratios for enrichment of population-specific var-CpGs in each genomic feature, separately for AFR, AMR, EAS, EUR and SAS compared with all other continental groups. Negative and positive values indicate depletion and enrichment, respectively. Bar colors denote continental groups. Red points indicate significant enrichment or depletion at FDR < 0.05, whereas gray points indicate nonsignificant associations. For visualization, $\log_2$ odds ratios were capped at −3 and 3. Txn, transcription; CNV, copy number variation.

**b**, Corresponding enrichment in pairwise continental group comparisons using $|\Delta| > 0.20$ and nominal $P < 5.0 \times 10^{-8}$.

**c**, Corresponding enrichment in pairwise continental group comparisons using $|\Delta| > 0.20$ and FDR < 0.05.

In **a–c**, the left and right panels show results for var-CpGs identified using the CpG gain/loss model and the integrated model combining CpG gain/loss with methylation, respectively. Colors indicate capped $\log_2$-transformed odds ratios, and black dots denote significant enrichment or depletion after Benjamini–Hochberg correction (FDR < 0.05).

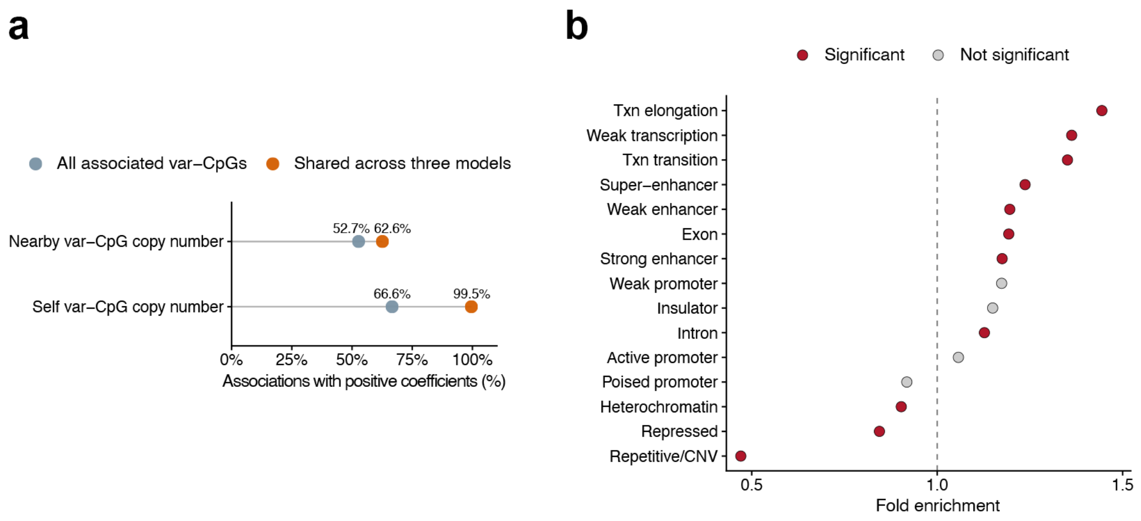


#### Extended Data Fig. 10: Directionality of methylation associations and genomic enrichment of shared var-CpGs.

**a**, Proportion of positive regression coefficients for associations between var-CpG methylation and either the copy number of the var-CpG itself or that of nearby var-CpGs. Results are shown for all associated var-CpGs in cyan blue and for the subset associated with all three genetic feature classes (shared var-CpGs) in orange. Shared var-CpGs showed pronounced positive directionality in associations with their own copy number.

**b**, Enrichment of shared var-CpGs across ChromHMM states and genomic features relative to the background set of var-CpGs whose methylation was associated with at least one genetic feature class. Each point represents one annotation category. The dashed vertical line indicates no enrichment (fold enrichment = 1). Red and gray points indicate significant and nonsignificant categories, respectively (FDR < 0.05). Enrichment was assessed using Fisher's exact tests followed by Benjamini–Hochberg correction. Txn, transcription; CNV, copy number variation.
