## Supplemental Data 2 for "Human panepigenome represents epigenomic diversity"

[*] *Corresponding author*

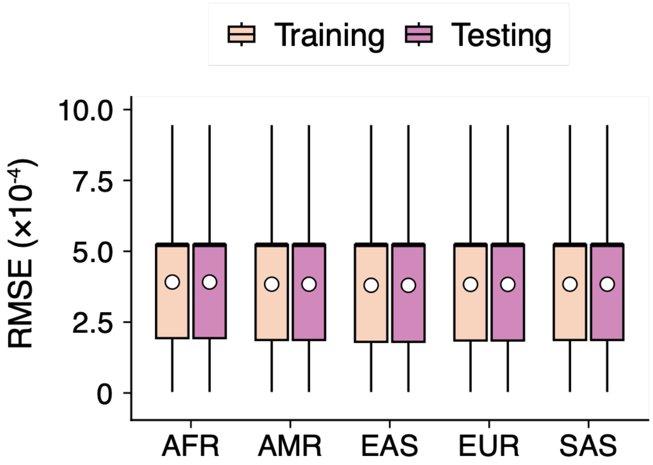


**Supplementary Fig. 1: Model prediction error across continental groups in the training and testing sets**.

Box plots show the distributions of root mean squared error (RMSE) between predicted and observed DNA methylation β-values, stratified by continental group and dataset. Boxes indicate the interquartile range, center lines denote medians, and whiskers extend to 1.5 times the interquartile range. White circles indicate means. Outliers are not displayed but were retained in the calculation of summary statistics.

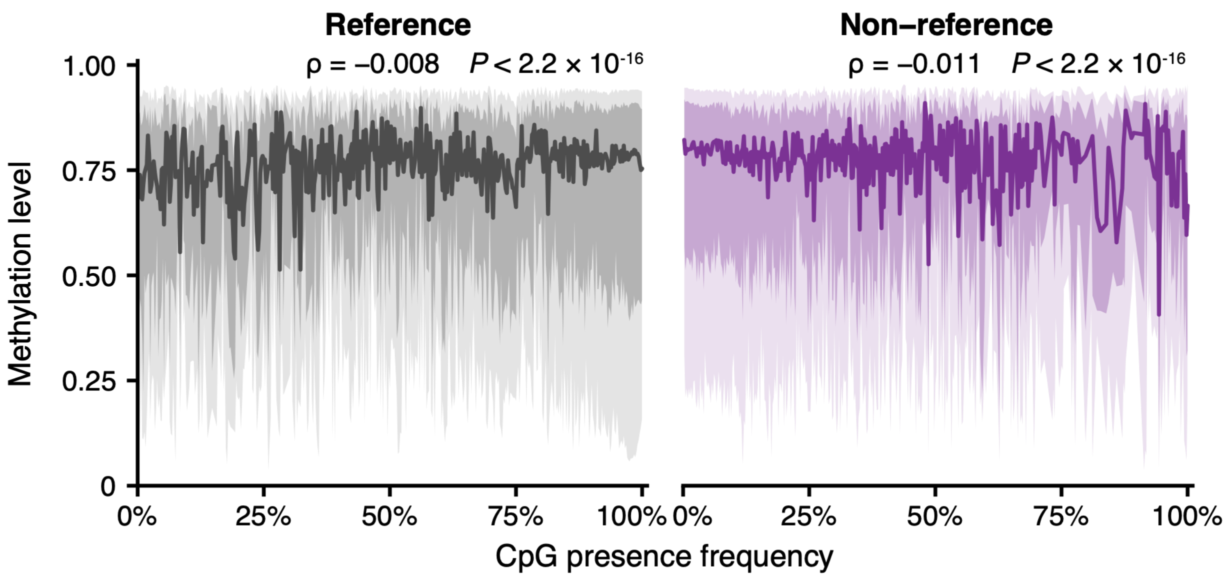


**Supplementary Fig. 2: Relationship between CpG presence frequency and methylation level.**

Solid curves show the median DNA methylation β-value across CpG presence frequencies in HPRC2, separately for reference and non-reference CpGs. Dark and light shaded regions indicate the interquartile range and the 10th–90th percentile range, respectively. For visualization, 1% of CpGs in each class were randomly sampled. Spearman's rank correlations between CpG presence frequency and methylation M-values were calculated separately for reference and non-reference CpGs using all CpGs. Spearman's ρ and two-sided *P* values are shown in each panel.
